# Intermediately Synchronised Brain States optimise trade-off between Subject Identifiability and Predictive Capacity

**DOI:** 10.1101/2022.09.30.510304

**Authors:** Leonard Sasse, Daouia I. Larabi, Amir Omidvarnia, Kyesam Jung, Felix Hoffstaedter, Gerhard Jocham, Simon B. Eickhoff, Kaustubh R. Patil

## Abstract

Functional connectivity (FC) refers to the statistical dependencies between activity of distinct brain areas. To study temporal fluctuations in FC within the duration of a functional magnetic resonance imaging (fMRI) scanning session, researchers have proposed the computation of an edge time series (ETS) and their derivatives. Evidence suggests that FC is driven by a few time points of high-amplitude co-fluctuation (HACF) in the ETS, which may also contribute disproportionately to interindividual differences. However, it remains unclear to what degree different time points actually contribute to brain-behaviour associations. Here, we systematically evaluate this question by assessing the predictive utility of FC estimates at different levels of co-fluctuation using machine learning (ML) approaches. We demonstrate that time points of lower and intermediate co-fluctuation levels provide overall highest subject specificity as well as highest predictive capacity of individual-level phenotypes.

## Introduction

In an effort to understand how brain organisation facilitates flexible yet specialised cognitive function, much neuroscientific research has focused on the functional connectivity (FC) between brain areas by investigating their pairwise correlations of functional magnetic resonance imaging (fMRI) blood oxygen level-dependent (BOLD) signals^1–3^. These pairwise correlations are assumed to represent the strength of connectivity (also called edges) between brain areas (also called nodes). Notably, considerable promise for the eventual applicability of FC as a biomarker has been shown with numerous studies demonstrating that FC differs between individuals, is stable within an individual^4–8^, and relates to individual-level cognition^4,9–13^ as well as clinically relevant symptoms of mental disorders^14–17^.

In order to move closer to the goal of applicability of FC biomarkers for real-world applications, researchers have tried to improve behavioural prediction by searching for the most suitable feature engineering schemes^9^, machine learning algorithms^10,12^, preprocessing parameters and brain parcellations^18,19^. Others focused on optimisation of within-individual stability and uniqueness of the fingerprint. This can be investigated within the identification framework in which the success of this optimisation is reflected in a higher success rate for the identification of an individual based on their FC profile or FC “fingerprint”^4^. With one such promising approach edge time series (ETS) are leveraged to select specific time points to estimate FC^20^. These time points of high-amplitude co-fluctuations, called “events”, contribute disproportionately to FC and are thought to reflect fluctuations in cognitive state^20^. These ETS reflect the magnitude of co-fluctuations of each pair of brain areas over time^21,22^ and are calculated as the product between the z-scored time series for each pair of brain areas. In order to study overall co-fluctuation patterns across all areas, the root sum of squares (RSS) at each time frame along the ETS has been suggested as a meaningful measure of co-fluctuation amplitude^20^. A higher RSS indicates higher co-fluctuations (or ETS) across the brain, i.e. higher overall brain synchronisation. This allows for the selection of only high-amplitude co-fluctuation (HACF) or low-amplitude co-fluctuation (LACF) time frames from the original BOLD time series. HACF-derived FC has been shown to yield enhanced subject identifiability, bringing up the question whether using these HACF time frames might also amplify brain-behaviour correlations^20,23,24^.

Thus far, a crucial missing component in the investigation of ETS is the evaluation of individual differences at different co-fluctuation amplitudes by means of prediction of phenotypes. Previous research has shown that connectivity of brain areas that contribute most to identification accuracy do not overlap with brain areas that contribute most to prediction accuracy^25^ suggesting that FC uniqueness and stability on their own do not guarantee phenotypic relevance of brain connectivity representations^26^. Furthermore, it is possible that different subsets of frames are more or less predictive of different phenotypic domains. In addition, one may ask, whether any selected subset of frames is more predictive than the full FC estimated using all available frames. Previous research has suggested that, to some degree, distinct brain states are differently associated to specific behaviours. For example, brain states showing strong integration between functional networks are associated with better cognitive task performance, in particular for memory- and attention-related tasks^27–29^. In addition, brain states characterised by high modularity are associated with better performance during motor tasks^30,31^.

Therefore, to address these questions, we systematically investigated the phenotypic relevance of FC contributions at different amplitudes of ETS co-fluctuations. Employing three time frame sampling strategies, we investigated predictiveness of targets across the domains of cognition, behaviour, personality, and demographics from FC including all frames, HACF frames, LACF frames, or a combination thereof using the Human Connectome Project (HCP-YA) S1200 dataset^32,33^. We also validate our findings using the Human Connectome Project Aging (HCP-A) dataset^34,35^. Further, to get a better understanding of what drives co-fluctuations, we investigated the influence of structural connectivity on the relationship between FC estimates at different co-fluctuation levels and predictiveness of these targets.

## Results

For each of the four resting state fMRI (rs-fMRI) runs of 771 subjects of the HCP-YA S1200 dataset^32,33^, we computed the edge time series (ETS) and the corresponding root sum of squares (RSS) along the ETS to quantify co-fluctuation amplitude using the Schaefer parcellation with 200 parcels^36^. For each subject, BOLD time series were ordered ranging from time frames with high overall co-fluctuation across the brain (high RSS; high amplitude co-fluctuation [HACF]) to low overall co-fluctuation (low amplitude co-fluctuation [LACF]). Next, we used three different strategies for selection of time frames with different levels of co-fluctuations: 1) sequential sampling: consecutively includes differing percentages of HACF or LACF using a threshold ranging from 0-50%, 2) individual bins: all time frames were divided into 20 individual bins each comprising 5% of time frames, and 3) combined bins: included time frames of all possible combinations of individual bins (see Methods for details; Fig. M1). FC matrices were created by calculating Pearson correlation coefficients between time series of each pair of brain areas while only including the selected time frames. Using these distinct FC matrices allowed us to systematically investigate the contributions of time frames with different levels of co-fluctuation to subject identifiability and prediction of 25 phenotypes.

### Differential identifiability and identification accuracy disagree in their assessment of FC “fingerprints”

In the sequential sampling strategy, we could replicate the finding that HACF frames yield higher differential identifiability (**I**_**Diff**_) than LACF frames (Fig. 1A). However, across the individual and combined bins sampling strategies it becomes evident that highest **I**_**Diff**_ is in fact achieved by intermediate bins (Fig. 1C). At the same time, the sequential sampling strategy shows that LACF frames provide higher identification accuracy (**I**_**Acc**_) than HACF frames (Fig. 1B). Yet, in the individual and combined bins sampling strategies, bins of intermediate co-fluctuation achieved the highest overall **I**_**Acc**_ (Fig. 1D). In the sequential sampling strategy, it is most apparent that **I**_**Acc**_ and **I**_**Diff**_ show opposing effects (Fig. 1A, B).

**Figure 1.**
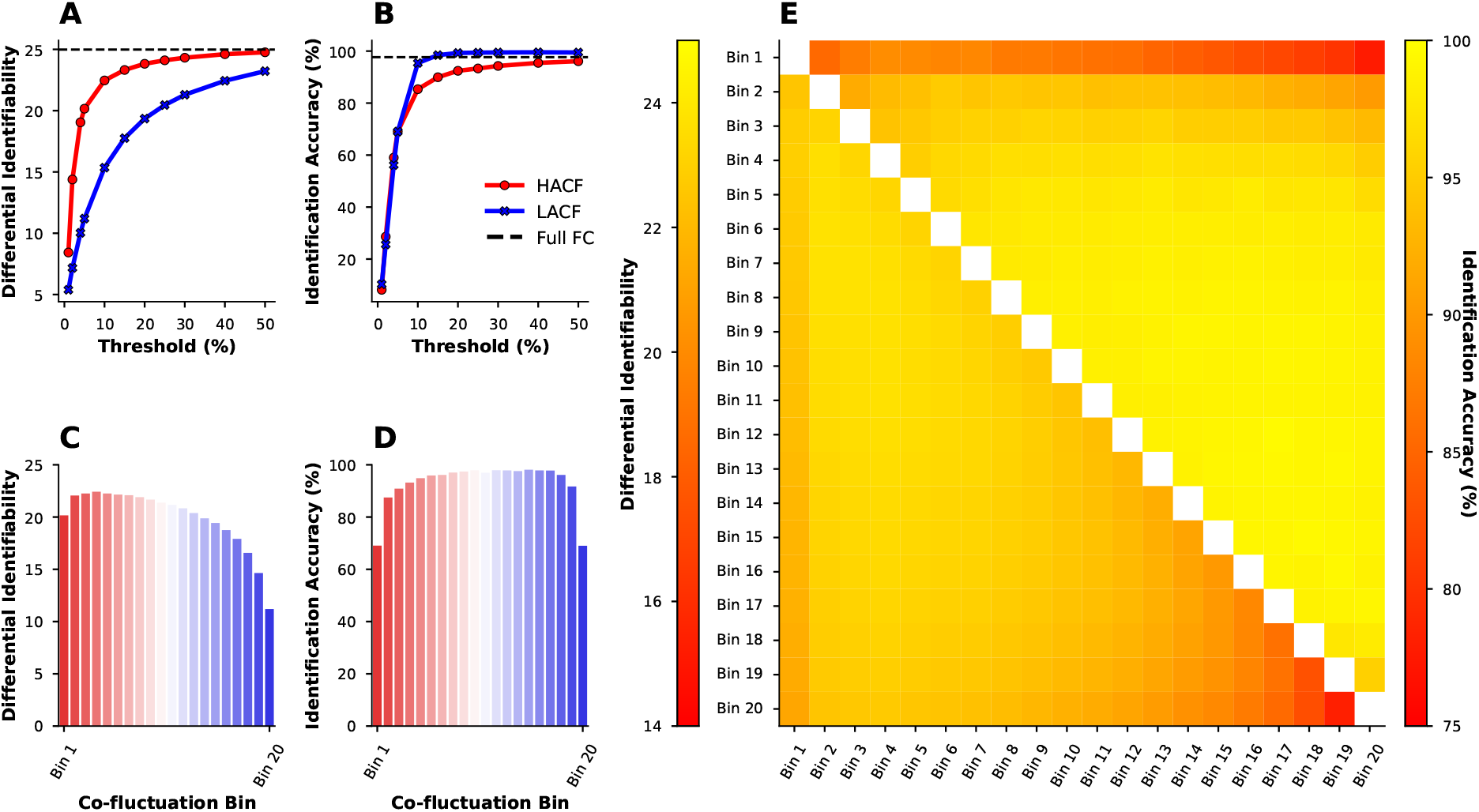
Differential identifiability and identification accuracy in the HCP-YA dataset for the sequential sampling strategy (**A** and **B**), the individual bins sampling strategy (**C** and **D**), and the combined bins strategy (**E**). In (**E**), the lower triangle shows differential identifiability, whereas the upper triangle shows identification accuracy achieved by each pair of combined bins.

### FC estimated at intermediate levels of co-fluctuation yield higher prediction accuracy than HACF or LACF frames

In order to test whether the differing behaviour of **I**_**Diff**_ and **I**_**Acc**_ can inform about the predictive utility, we then applied the sampling strategies to predict phenotypes using kernel ridge regression. We selected phenotypes used in ref.^37^ from the categories “Cognition”, “In-scanner task performance”, and “Personality” (see Table S1). Results displayed here consist of the 9 phenotypic targets with highest prediction accuracy based on the full FC. We report results for other difficult to predict targets (Figs. S4 and S5) as well as results using the coefficient of determination (*R*^2^) as scoring metric (Figs. S6 and S7) in the supplementary information. In the individual and combined bins sampling strategies a general trend can be observed with LACF frames yielding higher prediction scores than HACF frames (Fig. 2). Using a 5% Bayesian ROPE^38^(see Materials and Methods - Prediction of Behavioural and Demographic Measures) to compare each co-fluctuation bin’s performance against the performance of full FC, we found that across most targets co-fluctuation bins show predictive utility that is equivalent to the full FC (Fig. 2). However, in particular for the two targets that overall can be predicted with the highest accuracy (“Reading” and “Vocabulary”), it is apparent that HACF bins actually yield meaningfully lower prediction accuracy scores than full FC.

**Figure 2.**
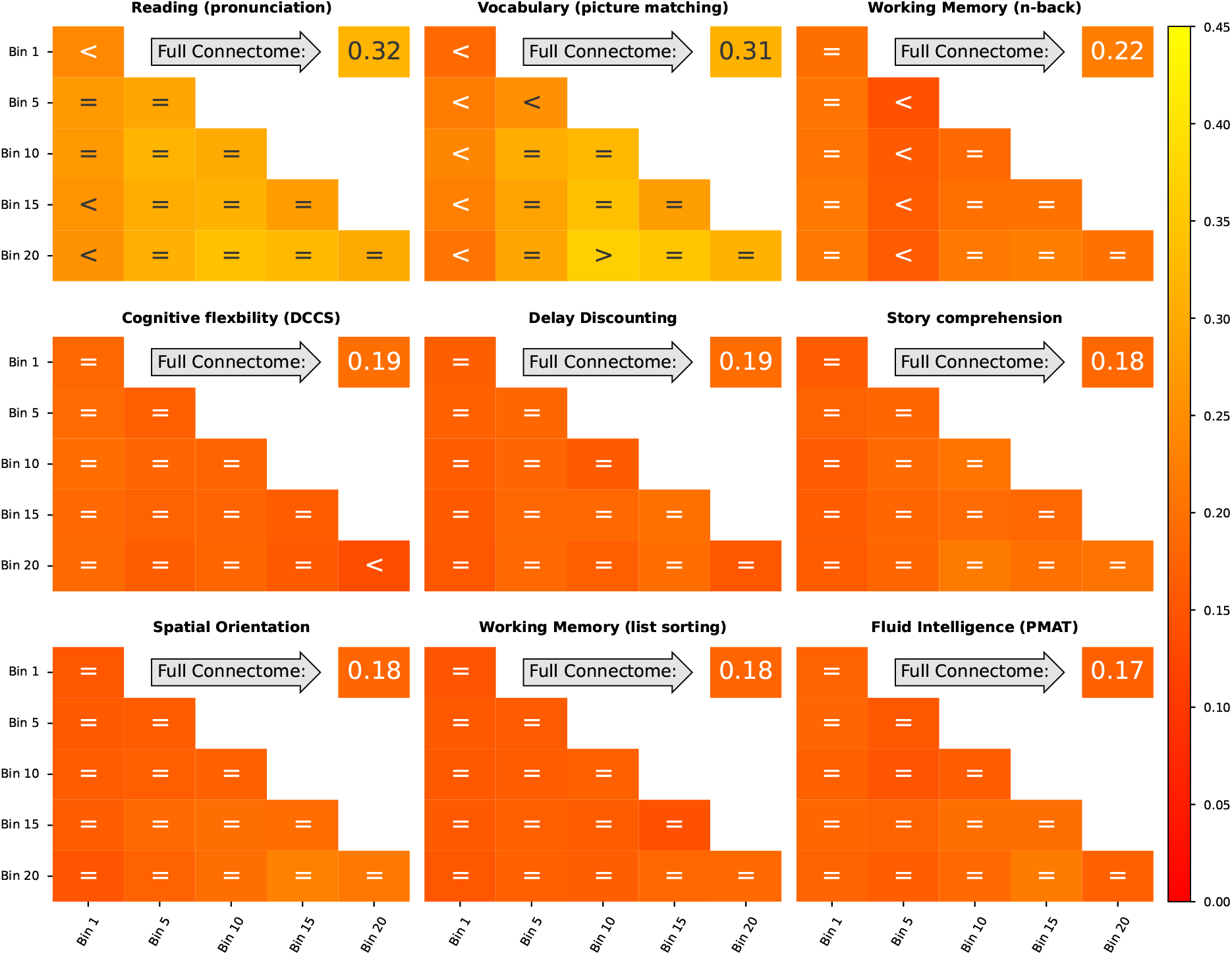
Prediction scores (Pearson’s r between observed and predicted values) for 9 phenotypic targets averaged across the ten folds in the grouped cross-validation scheme when using combined and individual bins sampling strategies. Bins range from 1 (HACF) to 20 (LACF). Scores for individual bins are displayed on the diagonal, for combined bins off the diagonal. Scores for the full FC using the whole time series are always displayed in the upper right corner. Comparison operators indicate whether scores obtained by a co-fluctuation bin are equivalent to scores obtained by full FC (“=“) or whether they are less (“<“) or greater (“>“) than scores obtained by full FC according to a 5% Bayesian ROPE^38^. These 9 targets are displayed, because they yielded best prediction accuracy using full FC compared to other targets displayed in the supplementary information.

In the sequential sampling strategy LACF frames also consistently yielded better prediction accuracy than HACF frames (Fig. 3). Results were consistent across scoring metrics. To illustrate robustness, we performed several additional analyses with different settings and parameters. To this end, we repeated the same analysis using Connectome-based Predictive Modelling (CBPM)^9^. Using this approach, prediction scores were lower, but overall a similar pattern was observed (see Figs. S8-S11 in the supplementary information). We further provide results for a subset of targets using the Schaefer 300 and 400 parcellations, with or without global signal regression, and for both scoring metrics, as these processing and analysis choices have been shown to affect prediction accuracy^19^ (Figs. S12-S21). For this analysis we chose “Reading” and “Vocabulary”, because they yielded best prediction accuracy scores as well as “Fluid Intelligence”, because it is widely used for prediction in the literature^4,39^.

**Figure 3.**
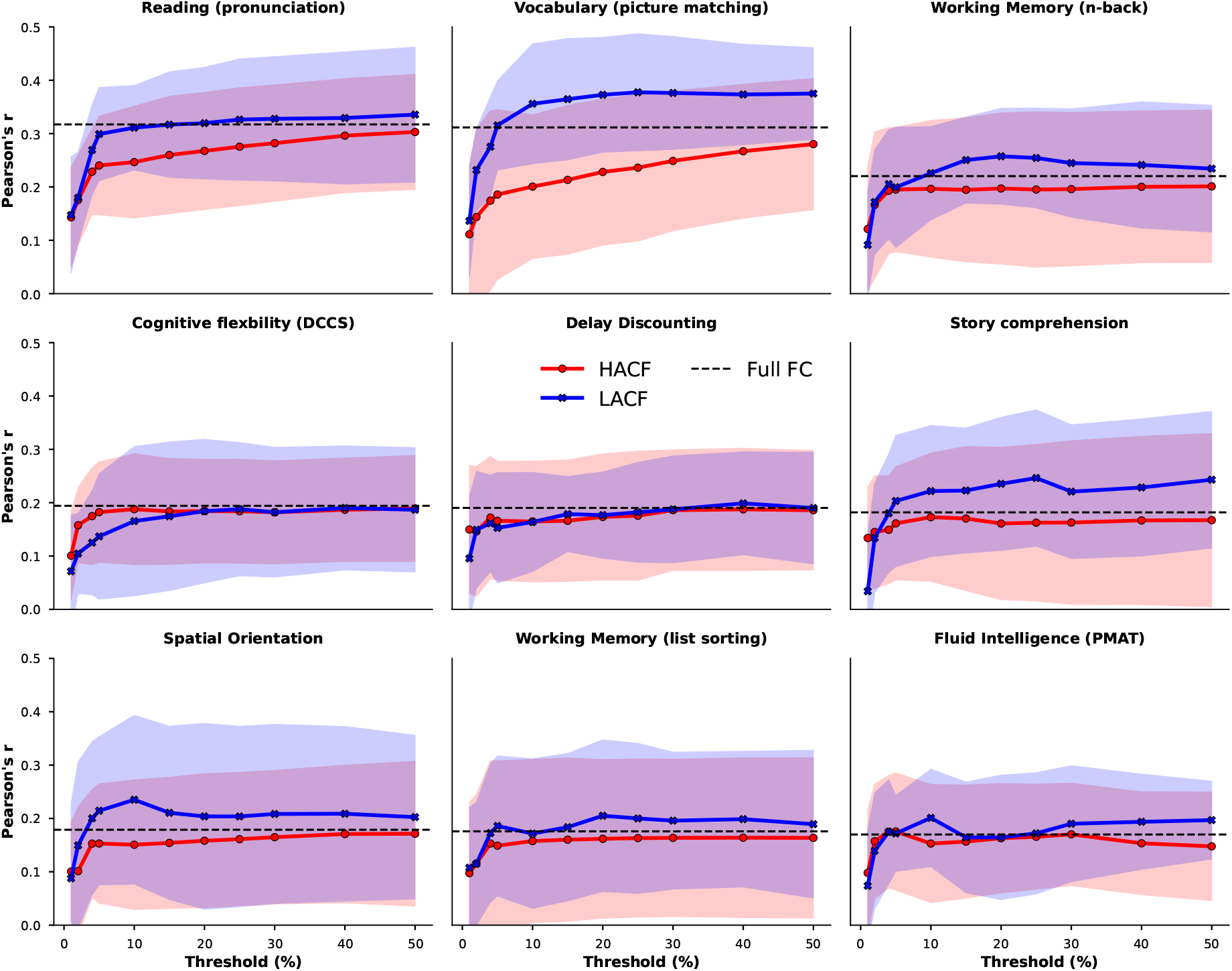
Prediction scores (Pearson’s r between observed and predicted) for 9 phenotypic targets averaged across the ten folds in the grouped cross-validation scheme when using FC estimates derived from time points at different levels of co-fluctuation magnitude using the sequential sampling strategy. Upper and lower boundary of the fill colours indicate the standard deviation across folds. A threshold of 100% corresponds to full FC. These 9 targets are displayed, because they yielded best prediction accuracy using full FC compared to other targets displayed in the supplementary information.

Overall, we made two robust observations across all analyses; LACF frames provide better predictive power than HACF frames, and some intermediate frames perform even better. To test whether these observations may be due to a relationship between the RSS and in-scanner motion, we correlated the RSS time series with framewise displacement (FD) for every subject and every rs-fMRI run. These correlations follow a normal distribution centered at zero and therefore show little evidence of a relationship between RSS and in-scanner motion (Fig. S22).

Given the importance of inter-individual demographic differences in basic and clinical research, we also tested whether this effect can be found when predicting age and sex which have shown better prediction accuracy than psychometric variables using FC^12,40^. In sex classification, we use a ridge classifier. In fact, a pattern similar to the previous analysis can be observed for both age and sex. In age prediction, the sequential sampling strategy clearly shows that HACF-derived FC yields lower prediction scores than LACF-derived FC (Fig. 4A). LACF-derived FC even yields higher prediction scores than the full FC, a pattern that can be found also in the combined bins strategy (Fig. 4E). In the individual bins strategy it is further evident that intermediate bins typically yield higher prediction scores than both HACF and LACF frames (Fig. 4C). We further report age prediction results using *R*^2^ (Fig. S23) and mean absolute error (MAE - Fig. S24) as scoring metrics. Considering the limited age range (22-37 years) in this sample, the age prediction scores are reasonable.

**Figure 4.**
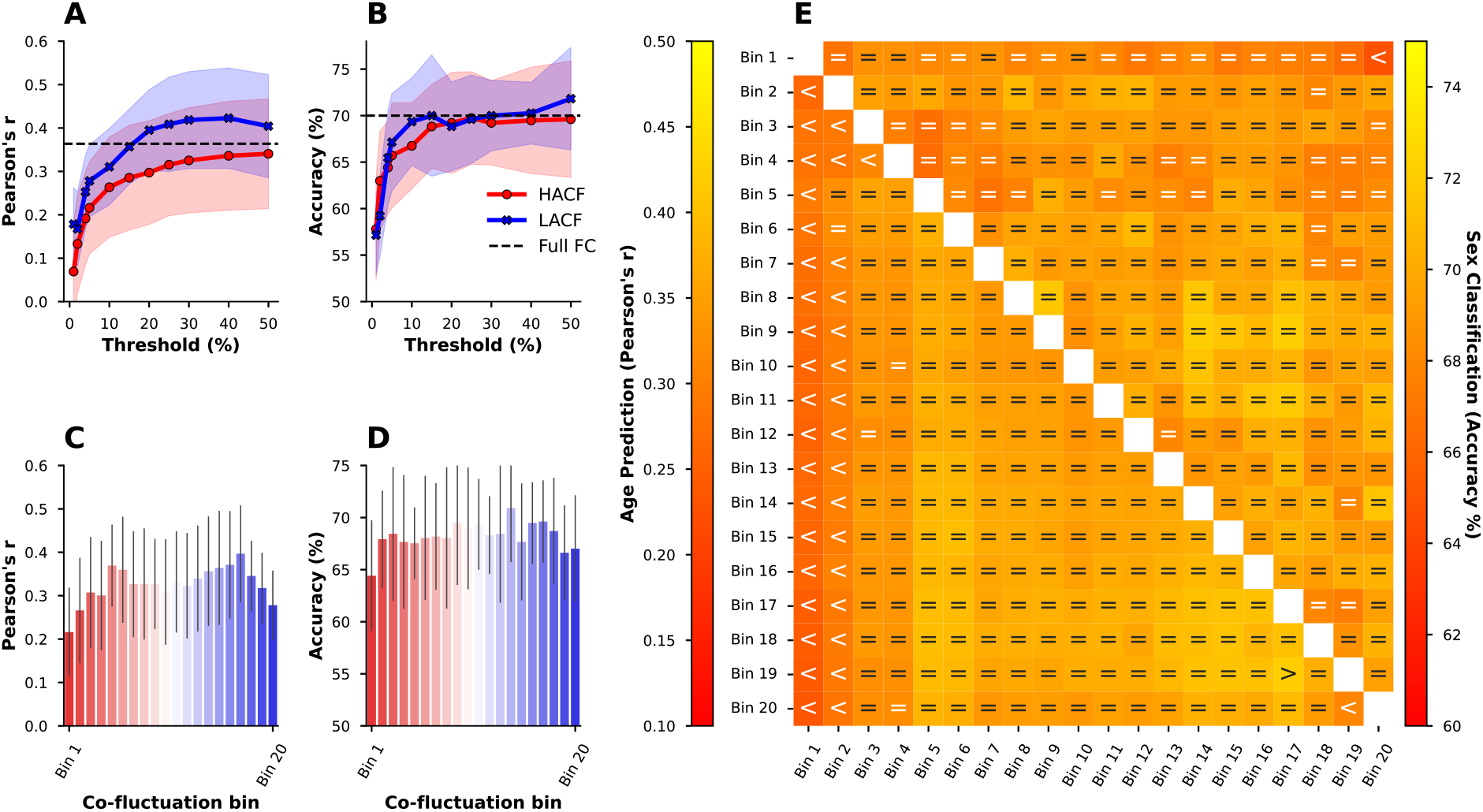
Age prediction (Pearson’s r) and sex classification (accuracy) using the combined bins (A), individual bins (B and D), and sequential increase (C and E) sampling strategies. Upper and lower boundary of the fill colours indicate the standard deviation across folds. Comparison operators indicate whether scores obtained by a co-fluctuation bin are equivalent to scores obtained by full FC (“=“) or whether they are less (“<“) or greater (“>“) than scores obtained by full FC according to a 5% Bayesian ROPE^38^

In sex classification, combined bins largely yielded prediction scores equivalent to full FC, however, a trend of prediction scores decreasing with higher co-fluctuation levels can be observed. In the individual bins strategy, both HACF and LACF bins yielded lower scores than intermediate bins. Lastly, comparing HACF and LACF frames using the sequential strategy, it can be seen that LACF frames usually provide better prediction scores than HACF frames. Again, to test whether results are robust across different models, we repeated sex prediction using support vector classifiers (SVC) with a linear kernel (Fig. S23) as well as a radial basis function (RBF) kernel (Fig. S24). The results were very similar.

### Validation dataset yields similar results

We then attempted to replicate identification and prediction in the HCP Aging (HCP-A) dataset (see Materials and Methods - Datasets). Results for **I**_**Acc**_ and **I**_**Diff**_ in the HCP-A dataset were consistent with the findings in the HCP-YA dataset across all sampling strategies (Fig. 5). **I**_**Acc**_ is higher for LACF frames than for HACF frames, whereas **I**_**Diff**_ is higher for HACF frames than for LACF frames, indicating that this effect is not sample-specific.

**Figure 5.**
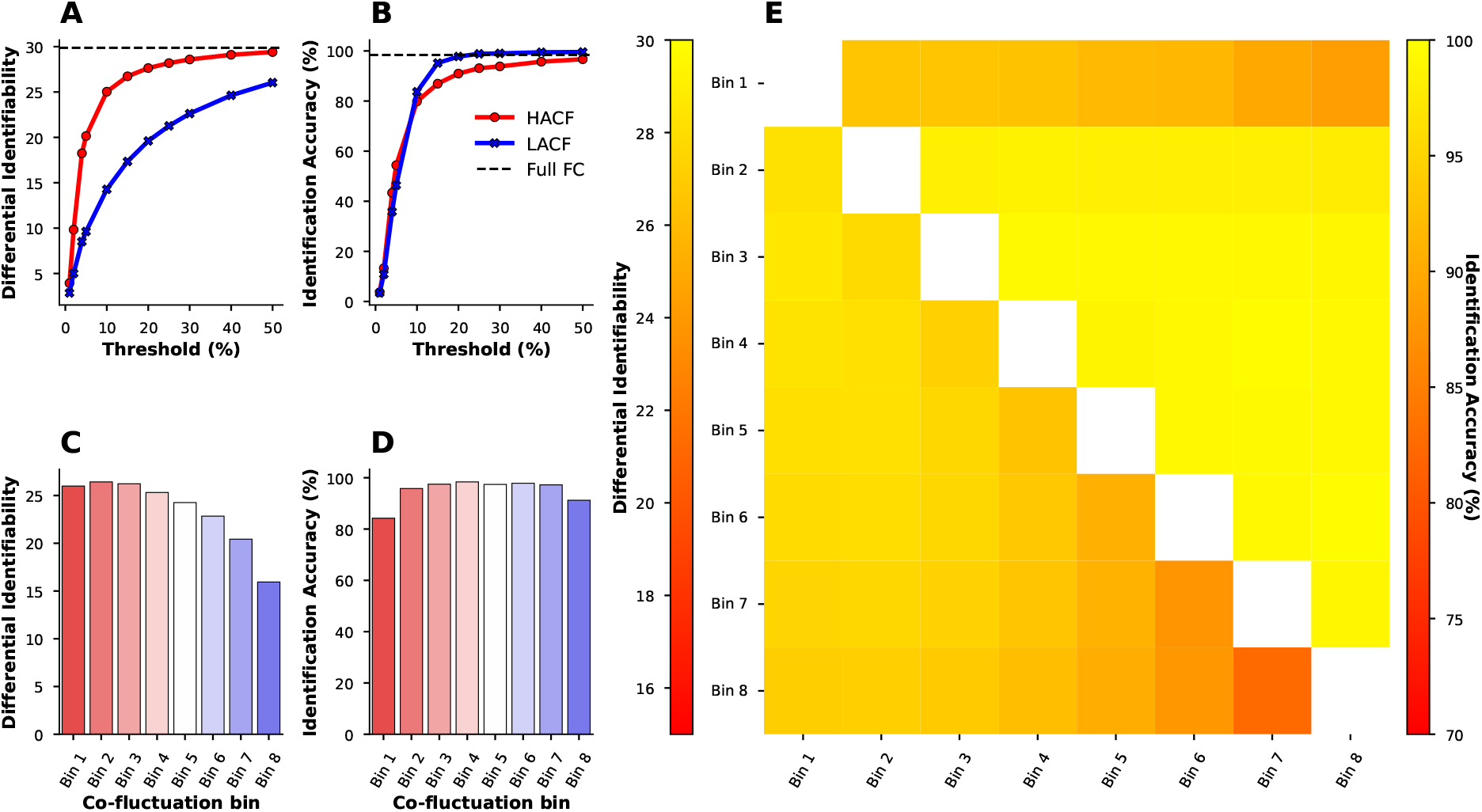
Differential identifiability and identification accuracy in the HCP-A dataset for the sequential sampling strategy (**A** and **B**), the individual bins sampling strategy (**C** and **D**), and the combined bins strategy (**E**).

For prediction in the HCP-A sample we selected “Language/Vocabulary Comprehension” and “Cognitive Flexibility” since these measures were also available in the HCP-YA sample and showed reasonable prediction accuracy. As there was no direct test of fluid intelligence in the HCP-A sample, we also included composite measures of fluid and crystallised cognition (see Table S2). Results on four cognitive targets show that prediction scores for these targets are overall largely equivalent to the full FC (Fig. 6A,B). However, a trend can be observed such that intermediate bins yield slightly higher performance than HACF or LACF frames. Similar results are obtained using *R*^2^ as scoring metric (Fig. S25). Next, we also set out to predict age and sex in the HCP-A dataset. However, since this sample was not balanced (316 female, 242 male), for sex classification we present balanced accuracy as scoring metric^41^. Results showed a similar pattern as previously observed but note the higher prediction accuracy for age on this sample due to its wider age range (Fig. 6C,E,G). In the individual and combined bins sampling strategies (Fig. 6E,F,G) scores are largely considered equivalent to prediction using the full FC, with a trend hinting at better performance in intermediate bins compared to HACF or LACF bins. Further, the sequential sampling strategy (Fig. 6C,D) makes it apparent that again LACF frames consistently yield better predictive performance compared to HACF frames. Again, in the supplementary material we report results for age prediction using *R*^2^ (Fig. S26) and MAE (Fig. S27) as well as results for sex prediction using a SVC with linear kernel (Fig. S26) and a RBF kernel (Fig. S27). Results for these parameters were consistent with the results displayed here.

**Figure 6.**
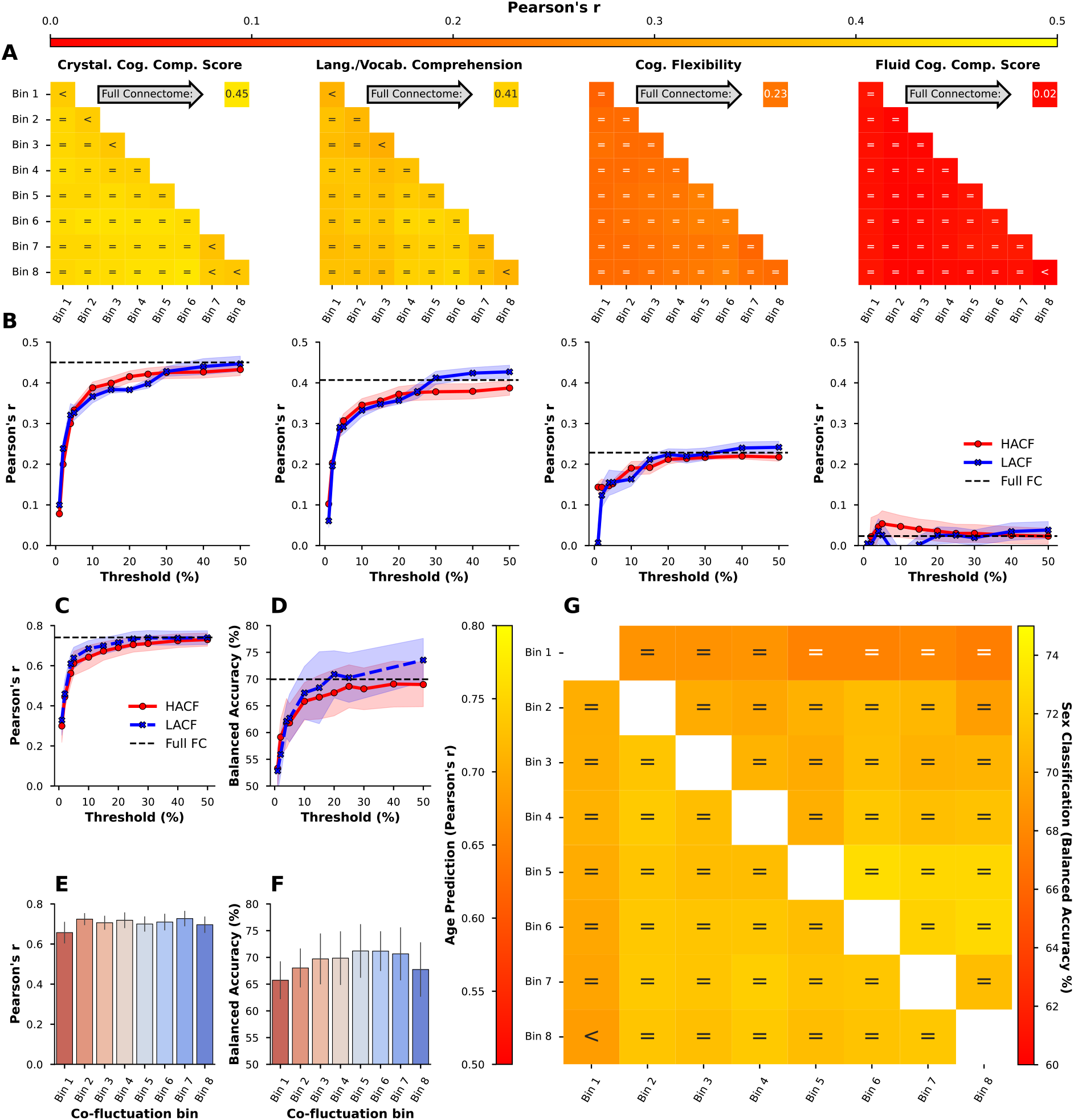
Prediction accuracy (Pearson’s r) for four cognitive targets in the HCP-A dataset using the individual and combined bins strategy (**A**; individual bins are on the diagonal) and the sequential sampling strategy (**B**). Upper and lower boundary of the fill colours indicate the standard deviation across repeats. Age prediction (Pearson’s r) and sex classification (balanced accuracy) using the sequential (**C** and **D**), individual bins (**E** and **F**), and combined bins (**G**) sampling strategies using the HCP-A sample. Comparison operators indicate whether scores obtained by a co-fluctuation bin are equivalent to scores obtained by full FC (“=“) or whether they are less (“<“) or greater (“>“) than scores obtained by full FC according to a 5% Bayesian ROPE^38^.

### FC shows a stronger Relationship to SC during intermediate Levels of Co-fluctuation

As a last analysis, we aimed to investigate the correspondence between FC at different levels of co-fluctuation and structural connectivity. As a first step, we correlated each subject’s SC with each co-fluctuation bin FC estimate. Our findings indicate that overall correlations between SC and FC tend to be greater for intermediate bins and some LACF bins compared to HACF bins (Fig. 7).

**Figure 7.**
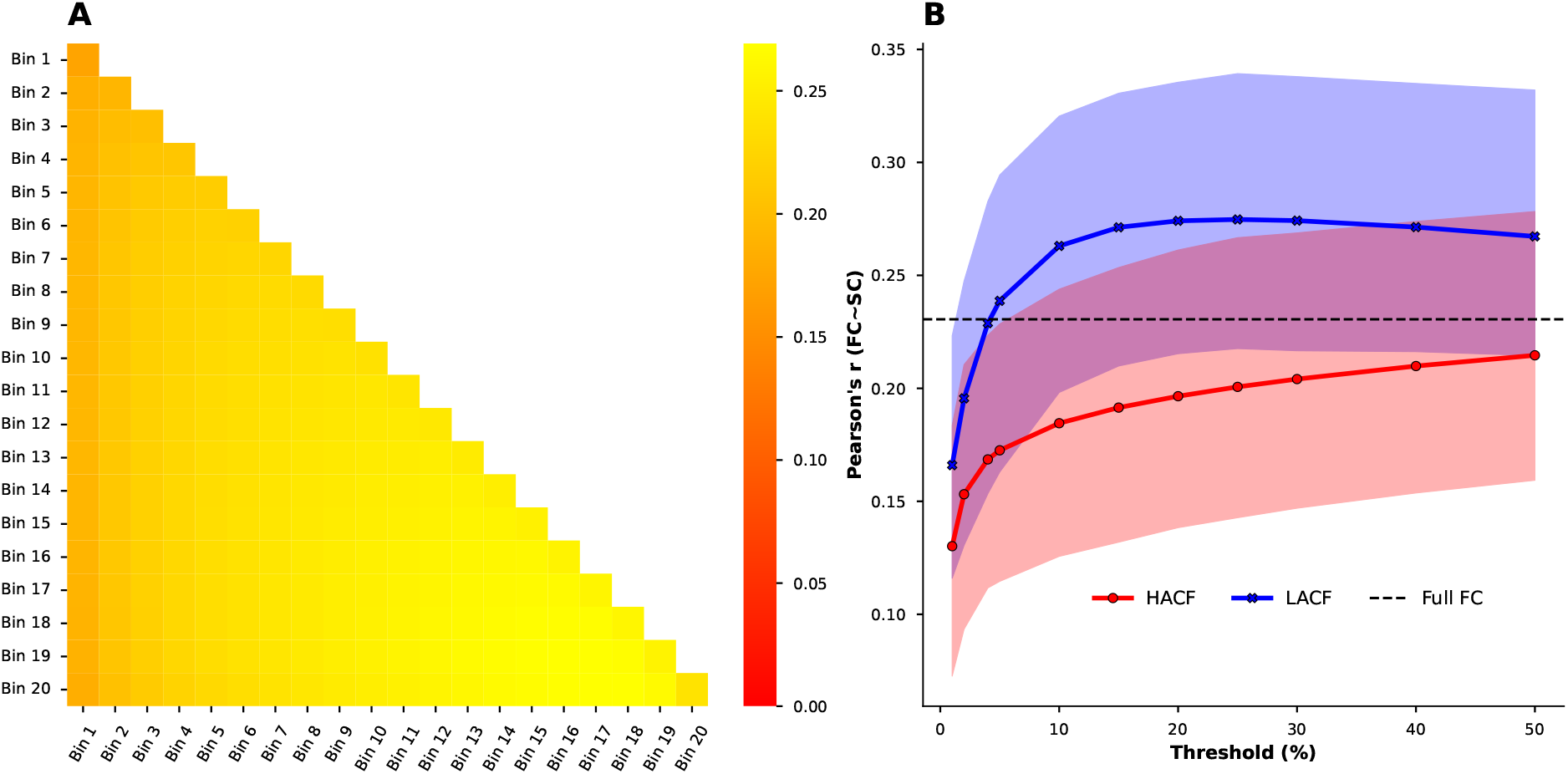
Correlations between SC and FC averaged across subjects in the **HCP-YA** sample for **A**) individual (on-diagonal) and combined bins (off-diagonal) and the **B**) sequential paradigm. In **B**) upper and lower bounds of fill colours indicate minimum and maximum correlations across subjects.

This trend is true for each of the three sampling strategies used. After obtaining these results we further wanted to test whether the information on underlying SC present in the intermediate bins and LACF bins indeed relates to the greater predictive utility of these bins (as compared to HACF bins). To test this we regressed out SC from the FC estimates for each subject and each co-fluctuation bin using a linear model and used the residuals for prediction of three cognitive variables in the HCP-YA dataset for which we observed a difference between HACF and LACF frames with respect to their predictive utility (Figs. 2, 3). The removal of SC from FC did not decidedly change prediction scores (Fig. 8A,B). We further repeated this paradigm in the prediction of sex and age. Here, the expected finding can be observed to a greater degree in the prediction of age (Fig. 8C, compare to Fig. 4A). Overall, however, regressing out SC does not seem to meaningfully change prediction scores, considering that the effect of intermediate frames and LACF frames yielding better predictions than HACF frames remains.

**Figure 8.**
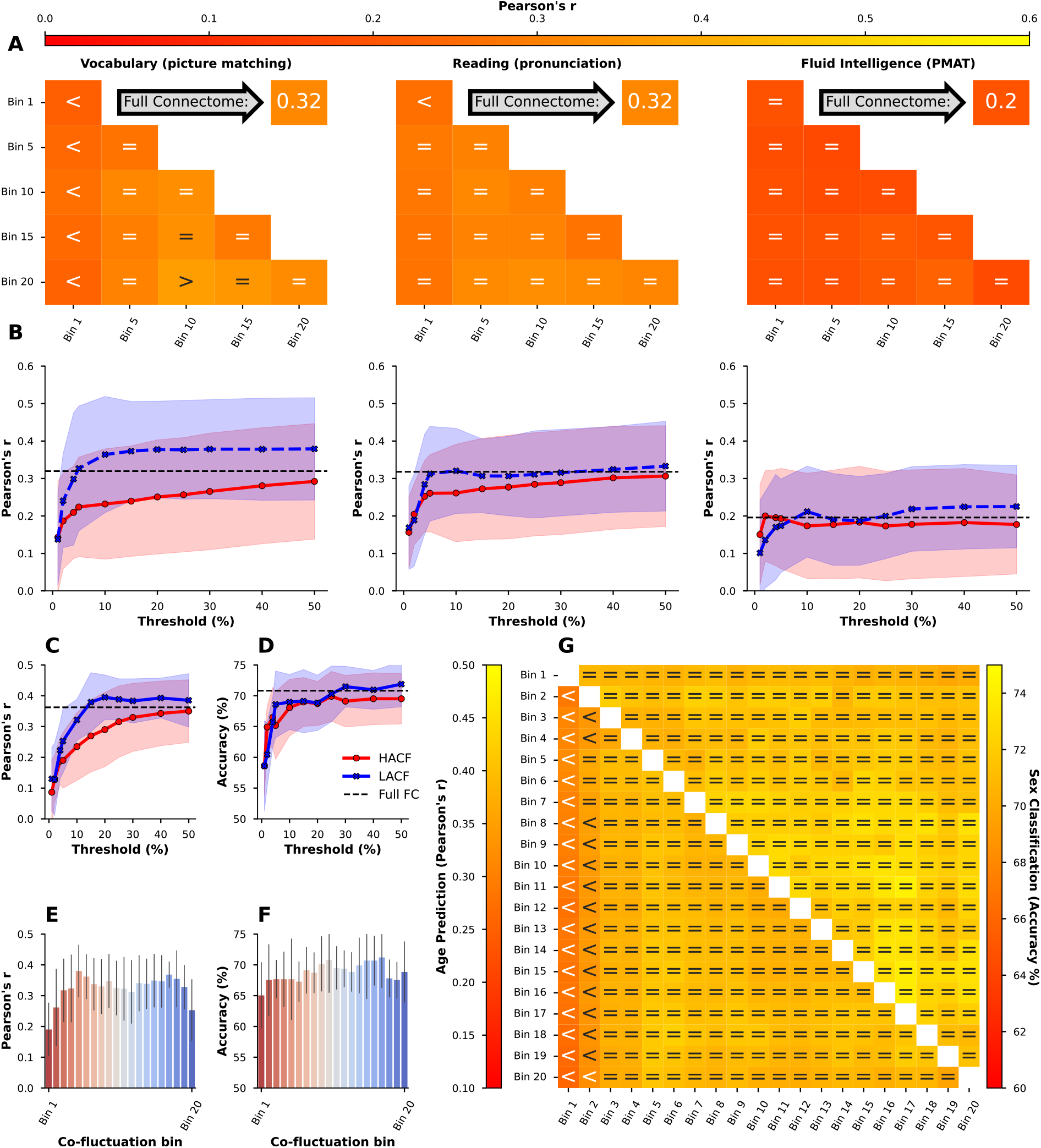
Prediction accuracy for three cognitive targets in the HCP-YA sample (Pearson’s r between observed and predicted targets) when removing SC from FC estimates for each subject in the combined and individual (**A**) and the sequential (**B**) sampling strategies. Age prediction (Pearson’s r) and sex classification (accuracy) using the sequential (**C** and **D**), individual bins (**E** and **F**), and combined bins (**G**) sampling strategies after removal of SC from FC. Comparison operators indicate whether scores obtained by a co-fluctuation bin are equivalent to scores obtained by full FC (“=“) or whether they are less (“<“) or greater (“>“) than scores obtained by full FC according to a 5% Bayesian ROPE^38^.

## Discussion

It has been suggested that the use of HACF frames with enhanced subject identifiability may amplify brain-behavior associations^24,42^. On the other hand, it has also been shown that FC-based identification and prediction may constitute conflicting goals^25,26^. Therefore, here we systematically evaluated the effect of inclusion of varying levels of functional co-fluctuations on subject identifiability and predictiveness of a range of phenotypes. Across a broad range of analytical settings and in two different cohorts, we observed that time frames with *intermediate* levels of co-fluctuation yield highest subject specificity (i.e. highest identification accuracies and differential identifiability between two rs-fMRI sessions) and greater predictive power compared to the other frames. Altogether, our findings suggest time frames with intermediate levels of co-fluctuation as a “sweet spot” capturing subject-specific phenotypic information; an intermediately synchronised and heterogeneous brain state situated in between a stereotypical highly synchronised brain state (HACF) and a weakly synchronised variable brain state (LACF). The highlighted role of intermediate co-fluctuation amplitudes in our study can be seen as a balanced point between increased synchrony (HACF) and increased disorder (LACF) between brain areas. Such a situation has also been reported at the brain neuronal level through the criticality hypothesis^43^ where neuronal activity of brain circuits self-organises into critical states or a transition between order and disorder. It also implies that the most relevant information of large-scale BOLD dynamics is encoded through discrete events in contrast to the dominant view of continuous fMRI data analysis^44^.

A previous study showed that differential identifiability of HACF-derived FC was higher than of LACF- derived FC^20^. Although we replicated this finding in our current study, we observed the opposite pattern for the identification problem with higher accuracies for LACF in contrast to HACF time frames. This finding is in line with another study on individual differences in FC estimated at different levels of co-fluctuation^42^, also showing that **I**_**diff**_ was highest in HACF frames while **I**_**acc**_ was highest at intermediate frames, and that removal of HACF frames decreased **I**_**diff**_ while increasing **I**_**Acc**_. Looking further into the subscores with which **I**_**diff**_ is calculated (within- and between-subject correlations), it can be seen that for HACF time frames the within-subject correlations increase more than the between-subject correlations (leading to higher **I**_**diff**_; see Figs. S1 and S2). This suggests a highly synchronised brain state that is stable within a subject. For LACF time frames, on the other hand, within-subject correlations decrease more than between-subject correlations (leading to lower **I**_**diff**_), suggesting this weakly synchronised brain state to be less stable within a subject. Integrating this with our identification accuracy findings, the lower identification accuracies for HACF suggest that this within-subject stable, highly synchronised brain state is also similar across subjects, i.e. a more “stereotypical” brain state^45^. Enhanced similarity between subjects at the HACF time frames is also in line with point process analysis findings where the spatial maps of resting state networks can be reconstructed from a limited number of high-amplitude events of fMRI^44^. It may explain higher uniformity of HACF time frames across the population. The higher identification accuracies for LACF may suggest that these weakly synchronised co-fluctuation amplitudes are less stable within a subject but even more variable across subjects.

Next, we systematically examined the predictiveness of different levels of co-fluctuation using machine learning based prediction analyses. In Zamani Esfahlani et al. (2020)^20^, an exploratory brain-behavior correlation analysis was performed. The authors performed PCA on 158 behavioral, trait, and demographic variables to obtain a first principal component (PC1) explaining 20.3% of behavioral variance. Correlations with top 5% HACF time frames as well as bottom 5% LACF time frames were weak, but relatively stronger for top 5% HACF time frames. In our study, we also found that the predictive capacity of FC estimates can indeed be improved by leveraging information from the ETS to select a specific set of points from the original BOLD time series. In contrast to Zamani Esfahlani et al. ^20^, we found lower predictiveness with HACF frames and higher predictiveness with LACF and especially intermediate time frames. This suggests that time points of low to intermediate levels of co-fluctuation may be best suited to make meaningful predictions. Using this insight in preprocessing pipelines may ultimately improve applicability of FC as an imaging-based biomarker of brain function in precision medicine and psychiatry.

A pivotal question associated with our findings concerns the origin of HACF and LACF events. Non-significant correlations have been shown between co-fluctuation amplitudes and breathing data, heart rate and motion (see Figs. S1 and S2 in ref.^20^), indicating that HACF events are unlikely to be driven by physiological noise only. Two main possibilities remain consistent with our results: On one hand there is evidence that HACF events are driven by tasks or change of cognitive state as demonstrated by the finding that RSS time series are more synchronised across participants during movie-watching than during resting state^20^. On the other hand it has been suggested that high RSS amplitudes are simply the result of extreme values in a noisy, but stationary distribution during rs-fMRI^46,47^. Temporal spacing of frames is likely an important factor to consider due to (temporal) autocorrelation^46^. Briefly, the reason that a few frames may suffice to faithfully reconstruct full FC, may be due to the fact that selected frames are well spaced along the time series, and therefore capture a broad range of points from that distribution. This may also play a relevant role in determining the predictive utility of specific FC estimates.

Assuming the validity of the first hypothesis, LACF and intermediate bins may then contain frames of brain activity during which estimated FC corresponds to a greater degree to a “baseline brain state” closer to underlying structural connectivity. To test this hypothesis, we investigated the correlation between FC at different levels of co-fluctuation and structural connectivity, measured by probabilistic tractography. Our findings indeed show higher correlations between FC during LACF and intermediate frames and structural connectivity (SC), just as another study showing that coupling between estimates of SC and FC was stronger during frames of intermediate or low levels of co-fluctuation^48^. This suggests that time resolved FC may inform us about how structural constraints drive functional organisation of the brain. In addition, language-related tasks (HCP Young adult: reading, vocabulary; HCP Aging: crystallized intelligence, language/vocabulary comprehension) were more predictable using intermediate and LACF time frames in contrast to HACF. This observation supports the hypothesis of a stronger link between SC and LACF time frames. It is also in line with the previous studies showing strong mapping between language and anatomy^49^. However, the predictive capacity of LACF and intermediate frames did not decrease after regressing out structural connectivity, suggesting these frames capture individual-level phenotypic information independent of brain structure (see Fig. 8). It seems unlikely therefore that the effect can only be explained by higher similarity with structural connectivity.

In light of these findings, it would be informative to investigate the properties of ETS in task-based fMRI (t-fMRI). If HACF events are in fact influenced by external stimuli, then one would expect more frequent HACF events after null model thresholding during task engagement. Otherwise, if the co-fluctuation levels and FC estimates over time do not reject stationary null models, then one could conclude that HACF events are simply the result of a random stationary process. Designing relevant null distributions for HACF/LACF of t-fMRI is an avenue for future work. A non-trivial challenge is due to the fact that computation of ETS requires z-scoring of the ROI time series, which is only appropriate if sample mean and standard deviation are time invariant^20,50^. A potential solution could be to regress out the block design from t-fMRI and use the residuals for this type of analysis.

### Limitations

One possible limitation when predicting behavioural variables is the unknown influence of confounding factors. In particular, the influence of in-scanner motion has been found to confound brain-behaviour relationships^51^. The impact of motion may also become influential in the identification and identifiability analysis through adding a highly personalized signature to the fMRI datasets^52^. However, as pointed out above, correlations between co-fluctuation amplitudes and physiological noise and motion have been found to be non-significant^20^. Further, the preprocessing strategies employed here to remove nuisance variables and influences of motion from the neural signal have been found to be among the most effective in the literature^53–55^. In addition, we also removed framewise displacement (FD; a measure of in-scanner head motion) from the predicted targets using a linear model. This was done in a CV-consistent fashion, meaning that a linear model was trained separately on the training data for each train-test split in the CV to avoid any test-to-train leakage of information regarding labels in the test set^56^. Lastly, we checked the correlations between RSS and FD for every subject and session, and found that correlations followed a normal distribution centered at zero (see Fig. S22).

Moreover, there may be other machine learning models not applied here, which perform better on FC estimates obtained during HACF frames than on FC estimates obtained during intermediate or LACF bins. However, in order to minimise this risk, we applied CBPM^4,9^ and kernel ridge regression, two models which are commonly used for FC-based prediction, and have been consistently found to yield competitive results in the prediction of cognitive and demographic variables^4,12,19,39,57^. Both models show results consistent with our conclusions here. Furthermore, we used three distinct models (linear SVM, RBF SVM and a ridge classifier) for sex classification, each of which confirmed the overall pattern that intermediate and LACF bins yield better prediction results than HACF bins.

### Conclusions and future research

It has previously been suggested that HACF time frames capture more individual-level information^20^. However, our findings suggest that time frames with intermediate levels of co-fluctuation yield highest subject identifiability and predictive capacity of individual-level phenotypes compared to HACF. We further find that assessments of subject identifiability provide more robust conclusions when multiple metrics are used (i.e. **I**_**diff**_ and **I**_**Acc**_). Overall, our findings suggest that intermediate frames may be more informative in individual-level inference and they may inform future preprocessing strategies aiming at identifying robust brain-based biomarkers.

## Materials and Methods

### Datasets - Human Connectome Project (HCP)

The details regarding collection of behavioural data, fMRI acquisition, and image preprocessing in the HCP Young-Adult (HCP-YA) project^32,33,58^ as well as the HCP Aging (HCP-A) project^34,35^ have been described elsewhere. Here, we aim to give a quick overview. The scanning protocol for both HCP-YA and HCP-A was approved by the local Institutional Review Board at Washington University in St. Louis. Retrospective analysis of these datasets was further approved by the local Ethics Committee at the Faculty of Medicine at Heinrich-Heine-University in Düsseldorf.

#### HCP Young-Adult (HCP-YA)

We used data obtained from two resting-state fMRI (rs-fMRI) sessions taken from the HCP-YA S1200 release^33^. Subjects were selected if data was available for both resting state sessions and 25 predefined behavioural variables of interest. This resulted in a dataset consisting of 771 subjects (384 female, 387 male). Participants’ age ranged from 22 to 37 (*M*=28.41, *SD*=3.74). Two sessions of rs-fMRI were obtained on two seperate days (each lasted ca. 15 minutes). Scans were acquired using a 3T Siemens connectome-Skyra scanner with a gradient-echo EPI sequence (TE=33.1ms, TR=720ms, flip angle = 52°, 2.0mm isotropic voxels, 72 slices, multiband factor of 8). For each fMRI session, scans were acquired for different phase encoding directions (left-right [LR] and right-left [RL]) providing four overall rs-fMRI datasets.

#### HCP Aging (HCP-A)

We used a subset of the HCP-A dataset as validation data to replicate our main findings from the exploratory HCP-YA dataset. Similar to the HCP-YA, in the HCP-A two sessions of rs-fMRI were acquired on two separate days with a 2D multiband (MB) gradient-recalled echo (GRE) echo-planar imaging (EPI) sequence (MB8, TR/TE = 800/37 ms, flip angle = 52°) and 2.0 mm isotropic voxels covering the whole brain (72 oblique-axial slices) using a Siemens 3T Prisma scanner. For each session, functional scans were acquired in two separate runs with opposite phase encoding polarity (anterior-to-posterior [AP] and posterior-to-anterior [PA]). Subjects who did not have data for all four of these runs were excluded. Further we only included subjects that had data for all four selected behavioural targets and confounding variables. This resulted in a sample of 558 subjects (316 female, 242 male) with ages ranging between 36 and 100 years (*M*=59.87, *SD*=15.03).

Data from rs-fMRI sessions in both of these datasets (HCP-YA and HCP-A) had also already undergone the HCP’s minimal preprocessing pipeline^32^, including motion correction and registration to standard space. Additionally, the ICA-FIX procedure (independent component analysis and FMRIB’s ICA-based X-noiseifier^54^) was applied to remove structured artefacts. Lastly, the 6 rigid-body parameters, their temporal derivatives and the squares of the 12 previous terms were regressed out, resulting in 24 parameters. Any further confound removal and preprocessing was applied to this denoised data.

### Image pre-processing

For both datasets, we regressed out confounds, linearly detrended and bandpass filtered the signal at 0.008 - 0.08 Hz using “nilearn.image.clean_img”. This included mean time courses of the white matter (WM), cerebro-spinal fluid (CSF), and global signal (GS), as well as their squared terms and temporal derivatives as confounds. A spike regressor was further added for each fMRI frame exceeding a motion threshold (0.25 mm root mean squared framewise displacement). The resulting voxel-wise images were then aggregated into the Schaefer 200 parcellation^36^. In the supplementary, we also provide results using the Schaefer 300 and 400 parcellation, and without the use of global signal regression. Additionally, in the HCP-A dataset time series were cut by excluding the first 20 and the last 18 volumes of the scan, so that the resulting time series consisted of 440 volumes that could be divided into 8 bins of 55 volumes. This was done to ensure that bins were of comparable size in both datasets, since the time series in the HCP-YA dataset (1200 volumes each) were divided into 20 bins of 60 volumes each.

### Edge Time Series Construction and FC Estimation

Edge time series were computed as described in previous research^20^. The parcellated BOLD time series were z-scored. Then, the element-wise product between the z-scored timeseries of each pair of parcels was computed as an estimate of co-fluctuation between ROIs over time. The magnitude of co-fluctuation was quantified using the root sum of squares at each timepoint (RSS) resulting in a co-fluctuation time series for each subject. Afterwards, the frames in the BOLD time series were ordered according to their corresponding RSS (from high to low) for every subject.

To test whether HACF moments capture more meaningful information about individual subjects than LACF moments, we sampled time frames using three separate strategies. For every subject and every resting-state session in the HCP dataset, the BOLD time series was ordered according to co-fluctuation magnitude. Each strategy differs in selection of frames used to construct the FC. In strategy 1) (individual bins sampling) the ranked BOLD time series was divided into twenty bins each containing 5% of the time series (60 frames) in the case of the HCP-YA dataset. From these twenty bins, five bins were sampled to be used in prediction. We did not consider all twenty bins in the HCP-YA in the prediction analysis, because running the pipeline for every bin and each of the targets would incur unnecessarily high and impractical computational cost. For each of the selected bins, FC was estimated using pairwise Pearson’s correlation coefficients. In the case of the HCP-A, the ranked BOLD time series was divided into 8 bins, each containing 12.5% of the timeseries (55 frames). All 8 bins were used in prediction. In strategy 2) (combined bins sampling), we sampled every possible combination of two bins out of the previously defined individual bins used in strategy 1) and concatenated these bins to estimate FC. In strategy 3) (sequential sampling), we used HACF and LACF frames, but applied sequentially increasing thresholds to include varying numbers of time points on either side. In each sampling strategy, FC estimates of corresponding co-fluctuation bins were first averaged across the phase encoding directions, resulting in two FC matrices per subject per co-fluctuation bin, to be used in identification. These two FC matrices were further averaged resulting in one FC matrix per subject per co-fluctuation bin to be used in prediction.

**Figure M1.**
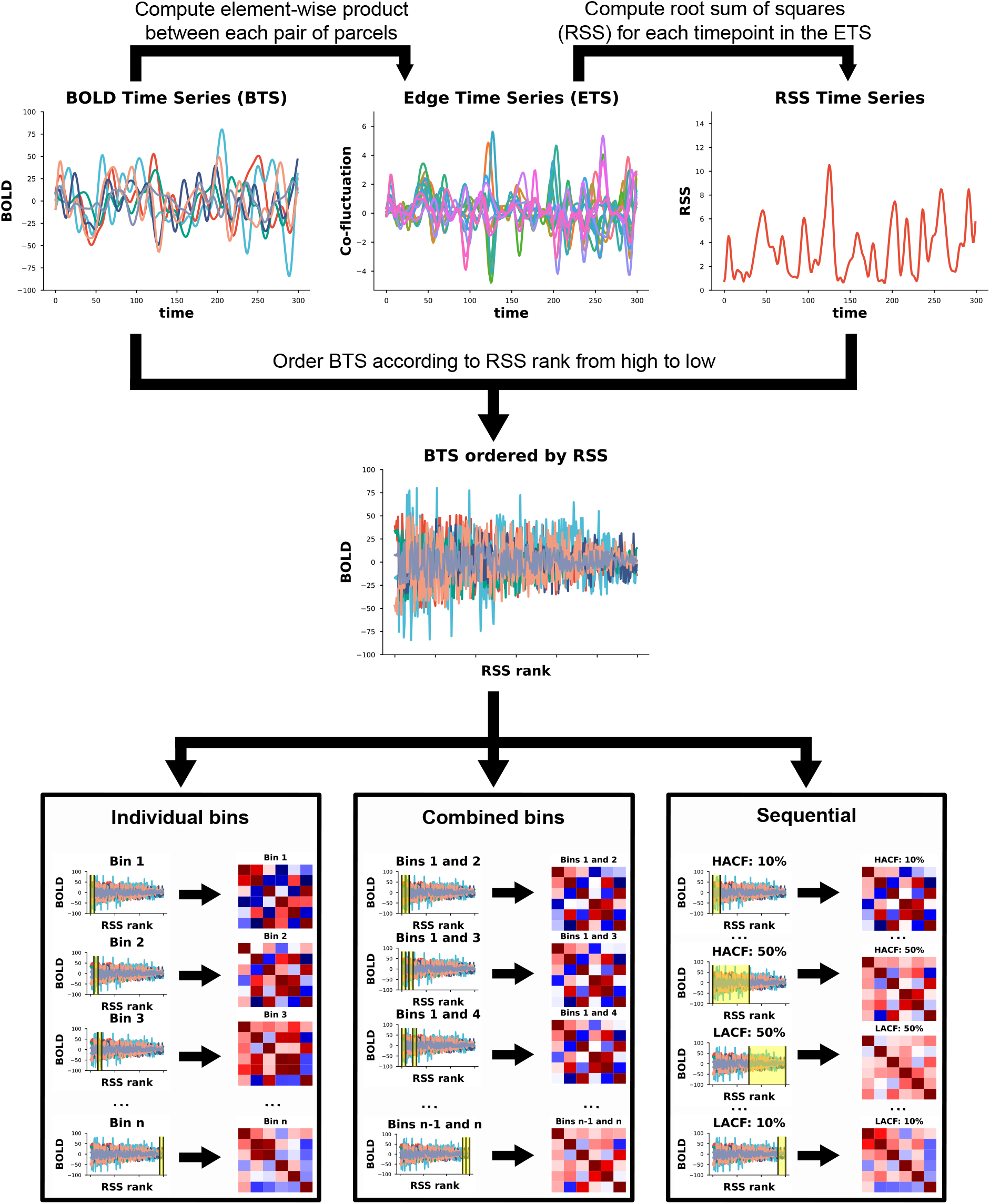
***Per subject workflow*** to extract FC at different levels of co-fluctuation using three different sampling strategies; **1)** in the ***individual bins strategy*** the re-ordered BOLD time series are divided into equally-sized bins, **2)**, in the ***combined bins strategy***, every possible combination of two bins was selected, frames from these two bins were concatenated, and **3)** in the ***sequential sampling strategy***, HACF or LACF frames were consecutively included to include only a percentage of the highest and lowest amplitude frames for each subject.

### Structural Connectivity Extraction

Diffusion-weighted magnetic resonance imaging (dMRI) data had already been processed using the HCP diffusion minimal preprocessing pipeline^32^. This included normalisation of the b0 image intensity across runs as well as removing EPI distortions, eddy-current-induced distortions and subject motion. It further corrected for gradient-nonlinearities and diffusion data was registered to the structural T1w scan. Structural connectivity (SC) matrices were extracted from these preprocessed dMRI data using a workflow developed in-house^59^. 10 million total streamlines of the whole-brain probabilistic tractography (WBT) were calculated using MRtrix3. Response functions were estimated using a 3-tissue constrained spherical deconvolution algorithm^60^. Fiber oriented distributions (FOD) were estimated from the dMRI data using spherical deconvolution, and the WBT was created through the fiber tracking by the second-order integration over the FOD by a probabilistic algorithm^61^. The tracking parameters were set as default values of the tckgen function from MRtrix documentation (https://mrtrix.readthedocs.io), where the following values were used: step size = 0.625 mm, angle = 45 degrees, minimal length = 2.5 mm, maximal length = 250 mm, FOD amplitude for terminating tract = 0.06, maximum attempts per seed = 1000, maximum number of sampling trials = 1000, and downsampling = 3. In the atlas transformation, labels were annotated using a classifier to parcel cortical regions in the native T1w space using Freesurfer^62^ according to the Schaefer atlas with 200-area parcellation^36^. The pipeline then transformed the labelled image from the T1w to dMRI native space. After the transformation, the labelled voxels in the grey matter mask were selected for a seed and a target region. Consequently, the “tck2connectome” function of MRtrix3 reconstructed SC. From the original sample of 771 subjects in the identification and prediction analyses, 3 subjects had to be excluded because they lacked dMRI data. Another 6 subjects were excluded due to software errors during structural preprocessing, resulting in a sample of 762 subjects (383 female, 379 male) with ages ranging from 22 to 37 (*M*=28.41, *SD*=3.73) for all analyses involving structural connectivity.

### Differential Identifiability and Identification Accuracy

To assess identifiability of FC for subjects across different sessions, we used both identification accuracy (**I**_**acc**_)^4^ and the differential identifiability quality function **I**_**diff**_^5^. “Identification” refers to the paradigm by which an individual’s FC profile obtained in an fMRI scanning session is used to identify them from a database of FC profiles obtained in a second fMRI scanning session^4^. While identification accuracy is defined as the proportion of correctly identified participants, differential identifiability is defined as the difference between mean within-subject correlations (**I**_**self**_) and mean between-subject correlations (**I**_**other**_): **I**_**diff**_ = (**I**_**self**_ *−* **I**_**other**_) *\** **100**^5,7^. Higher levels of differential identifiability indicate a stronger individual fingerprint.

### Prediction of Behavioural and Demographic Measures

Prediction was performed with FC matrices obtained using the three distinct strategies outlined above with unique edges serving as features. We selected phenotypes used in ref.^37^ from the categories “Cognition”, “In-scanner task performance”, and “Personality” (see Table S1) as targets, resulting in 25 targets overall. In the HCP-A dataset we used 4 cognitive targets. We selected “Language/Vocabulary Comprehension” and “Cognitive Flexibility” since these measures were also available in the HCP-YA sample and showed reasonable prediction accuracy. As there was no direct test of fluid intelligence in the HCP-A sample, we also included composite measures of fluid and crystallised cognition (see Table S2). Detailed descriptions of these targets can be found elsewhere^35,58^. To control for confounding influences, age, sex and framewise displacement (FD) were regressed out from the targets in a CV-consistent fashion as these have been found to correlate with behavioural variables^12,51^. In addition to prediction of behavioural targets we also predicted age and sex. In the prediction of age, only sex and FD were removed as confounds. In the prediction of sex, we removed age, brain volume (“FS_BrainSeg_Vol”), educational status (“SSAGA_Educ”), and FD as confounds. In the HCP-YA dataset, one subject (male) had to be excluded from sex prediction due to missing information on confounds (“SSAGA_Educ”).

For all regression tasks, we used ridge regression with a Pearson kernel. This model has been recommended as an efficient way to benchmark predictive utility of FC representations, and it performs well even compared to sophisticated deep learning algorithms specifically designed for connectivity-based features^12,18,19^. We further validated our main findings using CBPM^9^. In sex classification, we used a ridge classifier as well as support vector classifiers (SVC) with a linear kernel or a radial basis function (RBF) kernel, to see whether results are robust across different models.

To assess out-of-sample prediction accuracy in the HCP-YA dataset, a 10-fold nested cross-validation (CV) was performed for each FC representation. The folds were split such that family members were always within the same fold, so that independence between folds was maintained. To select the l2- regularisation strength for ridge regression and classification as well as the C parameter for the support vector classifiers in CV-consistent fashion, we used a 5-fold inner CV on the training folds. The model with the best parameters was fitted on the training folds and then tested on the outer CV test fold. In the HCP-A dataset we used a 5-fold nested CV with 5 repetitions, since we only included unrelated subjects, and therefore had no grouping constraint. Additionally, the number of samples was lower, so a 5-fold CV could ensure that test folds have sufficient samples. To evaluate prediction accuracy, we report Pearson’s r and the coefficient of determination (*R*^2^) for regression tasks as well as the mean absolute error (MAE) in the case of age prediction. For predicted values ŷ and corresponding observed values *y* over *n* samples with a sample mean of the observed values 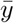, these metrics are defined as:

#### 1) Pearson’s r

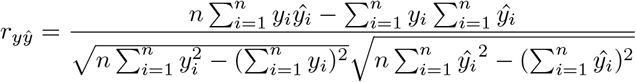

#### 2) Coefficient of Determination

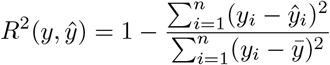

#### 3) MAE

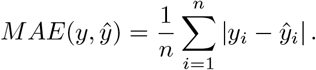

Unlike Pearson’s r and *R*^2^, the MAE is not scale invariant and therefore more difficult to interpret when predicting psychometric variables on different scales. In sex classification we used accuracy in the HCP-YA dataset:

#### 4) Accuracy

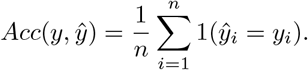

Due to imbalanced distribution of the class labels (316 female, 242 male), balanced accuracy was used in the HCP-A dataset to avoid inflated performance estimates:

#### 5) Balanced Accuracy

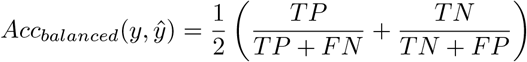

where *TP, FP, TN*, and *FN* denote true positives, false positives, true negatives, and false negatives respectively^41^.

To compare prediction accuracy between individual or combined bins and the full FC, we used a Bayesian ‘region of practical equivalence’ (ROPE) approach^38^. In this approach one defines two models as practically equivalent if differences between accuracy scores do not exceed a pre-defined percentage. Bayesian estimation^63^ can then be used to determine the probability that two models are practically equivalent and the probabilities that one model performs better than the other. One advantage of Bayesian estimation is that it does not rely on a point-wise null hypothesis but rather on a range of potential values (i.e. the prior distribution).

## Data and Code Availability

Further information on how to obtain the HCP-YA and the HCP-A datasets can be obtained at https://www.humanconnectome.org/. Code used to generate edge time series, connectomes, and perform prediction and other analyses on these data can be found in a public GitHub repository (https://github.com/juaml/etspredict). The code used to obtain SC is available at https://jugit.fz-juelich.de/inm7/public/vbcmri-pipeline.

Data and/or research tools used in the preparation of this manuscript were obtained from the National Institute of Mental Health (NIMH) Data Archive (NDA). NDA is a collaborative informatics system created by the National Institutes of Health to provide a national resource to support and accelerate research in mental health. Dataset identifier: [https://doi.org/10.15154/1527952]. This manuscript reflects the views of the authors and may not reflect the opinions or views of the NIH or of the Submitters submitting original data to NDA.

## Acknowledgements

This work was supported by the Helmholtz Portfolio Theme “Supercomputing and Modelling for the Human Brain” and the European Union’s Horizon 2020 Research and Innovation Programme under Grant Agreement No. 945539 (HBP SGA3).

## References

1. Biswal, B., Yetkin, F. Z., Haughton, V. M. & Hyde, J. S. Functional connectivity in the motor cortex of resting human brain using echo-planar MRI. Magnetic Resonance in Medicine 34, 537–541 (1995).

2. Craddock, R. C. et al. Imaging human connectomes at the macroscale. Nature Methods 10, 524–539 (2013).

3. Zalesky, A., Cocchi, L., Fornito, A., Murray, M. M. & Bullmore, E. Connectivity differences in brain networks. Neuroimage 60, 1055–1062 (2012).

4. Finn, E. S. et al. Functional connectome fingerprinting: Identifying individuals using patterns of brain connectivity. Nature Neuroscience 18, 1664–1671 (2015).

5. Amico, E. & Goñi, J. The quest for identifiability in human functional connectomes. Scientific Reports 8, 8254 (2018).

6. Bari, S., Amico, E., Vike, N., Talavage, T. M. & Goñi, J. Uncovering multi-site identifiability based on resting-state functional connectomes. Neuroimage 202, 115967 (2019).

7. Rajapandian, M., Amico, E., Abbas, K., Ventresca, M. & Goñi, J. Uncovering differential identifiability in network properties of human brain functional connectomes. Network neuroscience (Cambridge, Mass.) 4, 698–713 (2020).

8. Demeter, D. V. et al. Functional connectivity fingerprints at rest are similar across youths and adults and vary with genetic similarity. iScience 23, 100801 (2020).

9. Shen, X. et al. Using connectome-based predictive modeling to predict individual behavior from brain connectivity. Nature Protocols 12, 506–518 (2017).

10. Cui, Z. & Gong, G. The effect of machine learning regression algorithms and sample size on individualized behavioral prediction with functional connectivity features. Neuroimage 178, 622–637 (2018).

11. Sripada, C. et al. Prediction of neurocognition in youth from resting state fMRI. Molecular Psychiatry 25, 3413–3421 (2020).

12. He, T. et al. Deep neural networks and kernel regression achieve comparable accuracies for functional connectivity prediction of behavior and demographics. Neuroimage 206, 116276 (2020).

13. Chen, J. et al. Shared and unique brain network features predict cognitive, personality, and mental health scores in the ABCD study. Nature Communications 13, (2022).

14. Shannon, B. J. et al. Premotor functional connectivity predicts impulsivity in juvenile offenders. Proceedings of the National Academy of Sciences 108, 11241–11245 (2011).

15. Uddin, L. Q. et al. Salience networkBased classification and prediction of symptom severity in children with autism. JAMA Psychiatry 70, 869 (2013).

16. Lake, E. M. et al. The functional brain organization of an individual allows prediction of measures of social abilities transdiagnostically in autism and attention-deficit/hyperactivity disorder. Biological Psychiatry 86, 315–326 (2019).

17. Chen, J. et al. Intrinsic connectivity patterns of task-defined brain networks allow individual prediction of cognitive symptom dimension of schizophrenia and are linked to molecular architecture. Biological Psychiatry 89, 308–319 (2021).

18. Kong, R. et al. Individual-specific areal-level parcellations improve functional connectivity prediction of behavior. Cerebral Cortex 31, 4477–4500 (2021).

19. Li, J. et al. Global signal regression strengthens association between resting-state functional connectivity and behavior. Neuroimage 196, 126–141 (2019).

20. Zamani Esfahlani, F. et al. High-amplitude cofluctuations in cortical activity drive functional connectivity. Proceedings of the National Academy of Sciences of the United States of America 117, 28393–28401 (2020).

21. Faskowitz, J., Esfahlani, F. Z., Jo, Y., Sporns, O. & Betzel, R. F. Edge-centric functional network representations of human cerebral cortex reveal overlapping system-level architecture. Nature Neuroscience 23, 1644–1654 (2020).

22. Jo, Y. et al. The diversity and multiplexity of edge communities within and between brain systems. Cell Reports 37, 110032 (2021).

23. Betzel, R. F., Cutts, S. A., Greenwell, S., Faskowitz, J. & Sporns, O. Individualized event structure drives individual differences in whole-brain functional connectivity. Neuroimage 252, 118993 (2022).

24. Pope, M., Fukushima, M., Betzel, R. F. & Sporns, O. Modular origins of high-amplitude cofluctuations in fine-scale functional connectivity dynamics. Proceedings of the National Academy of Sciences of the United States of America 118, (2021).

25. Mantwill, M., Gell, M., Krohn, S. & Finke, C. Brain connectivity fingerprinting and behavioural prediction rest on distinct functional systems of the human connectome. Communications Biology 5, 261 (2022).

26. Finn, E. S. & Rosenberg, M. D. Beyond fingerprinting: Choosing predictive connectomes over reliable connectomes. Neuroimage 239, 118254 (2021).

27. Braun, U. et al. Dynamic reconfiguration of frontal brain networks during executive cognition in humans. Proceedings of the National Academy of Sciences of the United States of America 112, 11678–11683 (2015).

28. Shine, J. M. et al. The dynamics of functional brain networks: Integrated network states during cognitive task performance. Neuron 92, 544–554 (2016).

29. Spadone, S. et al. Dynamic reorganization of human resting-state networks during visuospatial attention. Proceedings of the National Academy of Sciences of the United States of America 112, 8112–8117 (2015).

30. Bassett, D. S., Yang, M., Wymbs, N. F. & Grafton, S. T. Learning-induced autonomy of sensorimotor systems. Nature Neuroscience 18, 744–751 (2015).

31. Mohr, H. et al. Integration and segregation of large-scale brain networks during short-term task automatization. Nature Communications 7, 13217 (2016).

32. Glasser, M. F. et al. The minimal preprocessing pipelines for the human connectome project. Neuroimage 80, 105–124 (2013).

33. Van Essen, D. C. et al. The WU-minn human connectome project: An overview. Neuroimage 80, 62–79 (2013).

34. Harms, M. P. et al. Extending the human connectome project across ages: Imaging protocols for the lifespan development and aging projects. Neuroimage 183, 972–984 (2018).

35. Bookheimer, S. Y. et al. The lifespan human connectome project in aging: An overview. Neuroimage 185, 335–348 (2019).

36. Schaefer, A. et al. Local-global parcellation of the human cerebral cortex from intrinsic functional connectivity MRI. Cerebral Cortex 28, 3095–3114 (2018).

37. Kong, R. et al. Spatial topography of individual-specific cortical networks predicts human cognition, personality, and emotion. Cerebral Cortex 29, 2533–2551 (2019).

38. Benavoli, A., Corani, G., Demšar, J. & Zaffalon, M. Time for a change: A tutorial for comparing multiple classifiers through bayesian analysis. Journal of Machine Learning Research (2017).

39. Vieira, B. H. et al. On the prediction of human intelligence from neuroimaging: A systematic review of methods and reporting. Intelligence 93, 101654 (2022).

40. Weis, S. et al. Sex classification by resting state brain connectivity. Cerebral Cortex 30, 824–835 (2020).

41. Brodersen, K. H., Ong, C. S., Stephan, K. E. & Buhmann, J. M. The balanced accuracy and its posterior distribution. In 2010 20th international conference on pattern recognition 3121–3124 (IEEE, 2010).

42. Cutts, S. A., Faskowitz, J., Betzel, R. F. & Sporns, O. Uncovering individual differences in fine-scale dynamics of functional connectivity. Cerebral Cortex (2022).

43. Zimmern, V. Why brain criticality is clinically relevant: A scoping review. Frontiers in Neural Circuits 14, (2020).

44. Tagliazucchi, E., Balenzuela, P., Fraiman, D. & Chialvo, D. R. Criticality in large-scale brain FMRI dynamics unveiled by a novel point process analysis. Frontiers in physiology 3, 15 (2012).

45. Betzel, R. F. et al. Hierarchical organization of spontaneous co-fluctuations in densely-sampled individuals using fMRI. BioRxiv (2022).

46. Ladwig, Z. et al. BOLD cofluctuation ‘events’ are predicted from static functional connectivity. NeuroImage 260, 119476 (2022).

47. Novelli, L. & Razi, A. A mathematical perspective on edge-centric brain functional connectivity. Nature Communications 13, 2693 (2022).

48. Liu, Z.-Q. et al. Time-resolved structure-function coupling in brain networks. Communications Biology 5, 532 (2022).

49. Dhamala, E., Jamison, K. W., Jaywant, A., Dennis, S. & Kuceyeski, A. Distinct functional and structural connections predict crystallised and fluid cognition in healthy adults. Human Brain Mapping 42, 3102–3118 (2021).

50. Liégeois, R., Laumann, T. O., Snyder, A. Z., Zhou, J. & Yeo, B. T. T. Interpreting temporal fluctuations in resting-state functional connectivity MRI. Neuroimage 163, 437–455 (2017).

51. Siegel, J. S. et al. Data quality influences observed links between functional connectivity and behavior. Cerebral Cortex 27, 4492–4502 (2017).

52. Bolton, T. A. et al. Agito ergo sum: Correlates of spatio-temporal motion characteristics during fMRI. NeuroImage 209, 116433 (2020).

53. Xifra-Porxas, A., Kassinopoulos, M. & Mitsis, G. D. Physiological and motion signatures in static and time-varying functional connectivity and their subject identifiability. eLife 10, (2021).

54. Salimi-Khorshidi, G. et al. Automatic denoising of functional MRI data: Combining independent component analysis and hierarchical fusion of classifiers. Neuroimage 90, 449–468 (2014).

55. Parkes, L., Fulcher, B., Yücel, M. & Fornito, A. An evaluation of the efficacy, reliability, and sensitivity of motion correction strategies for resting-state functional MRI. Neuroimage 171, 415–436 (2018).

56. More, S., Eickhoff, S. B., Caspers, J. & Patil, K. R. Confound removal and normalization in practice: A neuroimaging based sex prediction case study. In ECML PKDD 2020: Demo track (eds. Dong, Y., Ifrim, G., Mladenic, D., Saunders, C. & Van Hoecke, S.) vol. 12461 3–18 (Springer International Publishing, 2021).

57. Dadi, K. et al. Benchmarking functional connectome-based predictive models for resting-state fMRI. Neuroimage 192, 115–134 (2019).

58. Barch, D. M. et al. Function in the human connectome: Task-fMRI and individual differences in behavior. Neuroimage 80, 169–189 (2013).

59. Jung, K., Eickhoff, S. B. & Popovych, O. V. Tractography density affects whole-brain structural architecture and resting-state dynamical modeling. Neuroimage 237, 118176 (2021).

60. Dhollander, T., Mito, R., Raffelt, D. & Connelly, A. Improved white matter response function estimation for 3-tissue constrained spherical deconvolution. ISMRM (2019).

61. Tournier, J., Calamante, F. & Connelly, A. Improved probabilistic streamlines tractography by 2nd order integration over fibre orientation distributions. ISMRM (2010).

62. Dale, A., Fischl, B. & Sereno, M. Cortical surface-based analysis. I. Segmentation and surface reconstruction. Neuroimage 9, 179–194 (1999).

63. Gelman, A. et al. Bayesian data analysis. (Chapman; Hall/CRC, 2013).

