## Supplementary Information for "Intermediately Synchronised Brain States optimise trade-off between Subject Identifiability and Predictive Capacity"

September 30, 2022

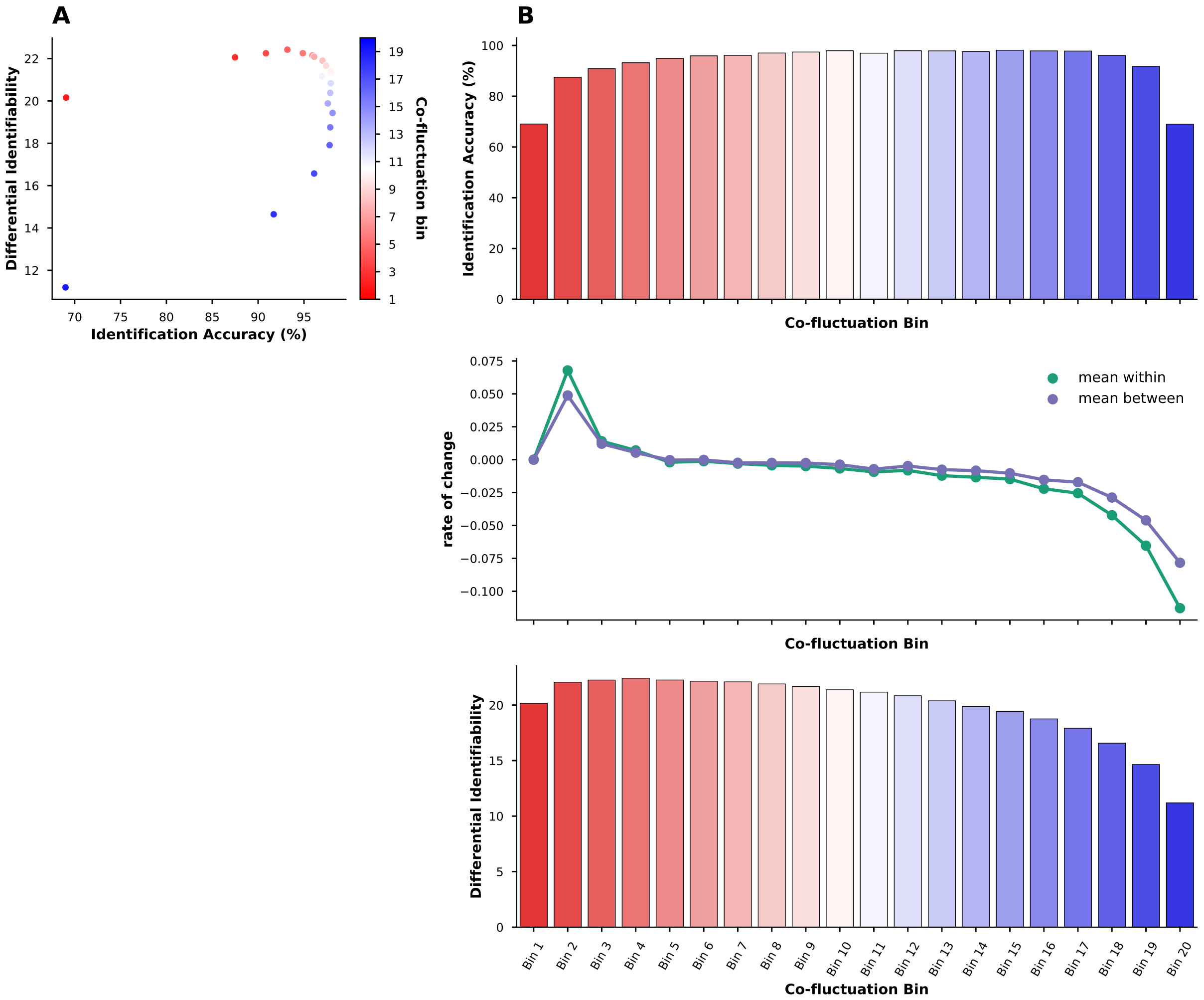


**Figure S1.** Identification accuracy and differential identifiability in the HCP-YA sample using the individual bins strategy: **A)** shows optimal subject specificity of intermediate bins in a scatterplot of identification accuracy and differential identifiability. **B** shows identification accuracy (top row) and differential identifiability (bottom row) as well as the derivative (i.e. difference between one bin and the next) of mean within- and mean-between subject correlations (middle row).

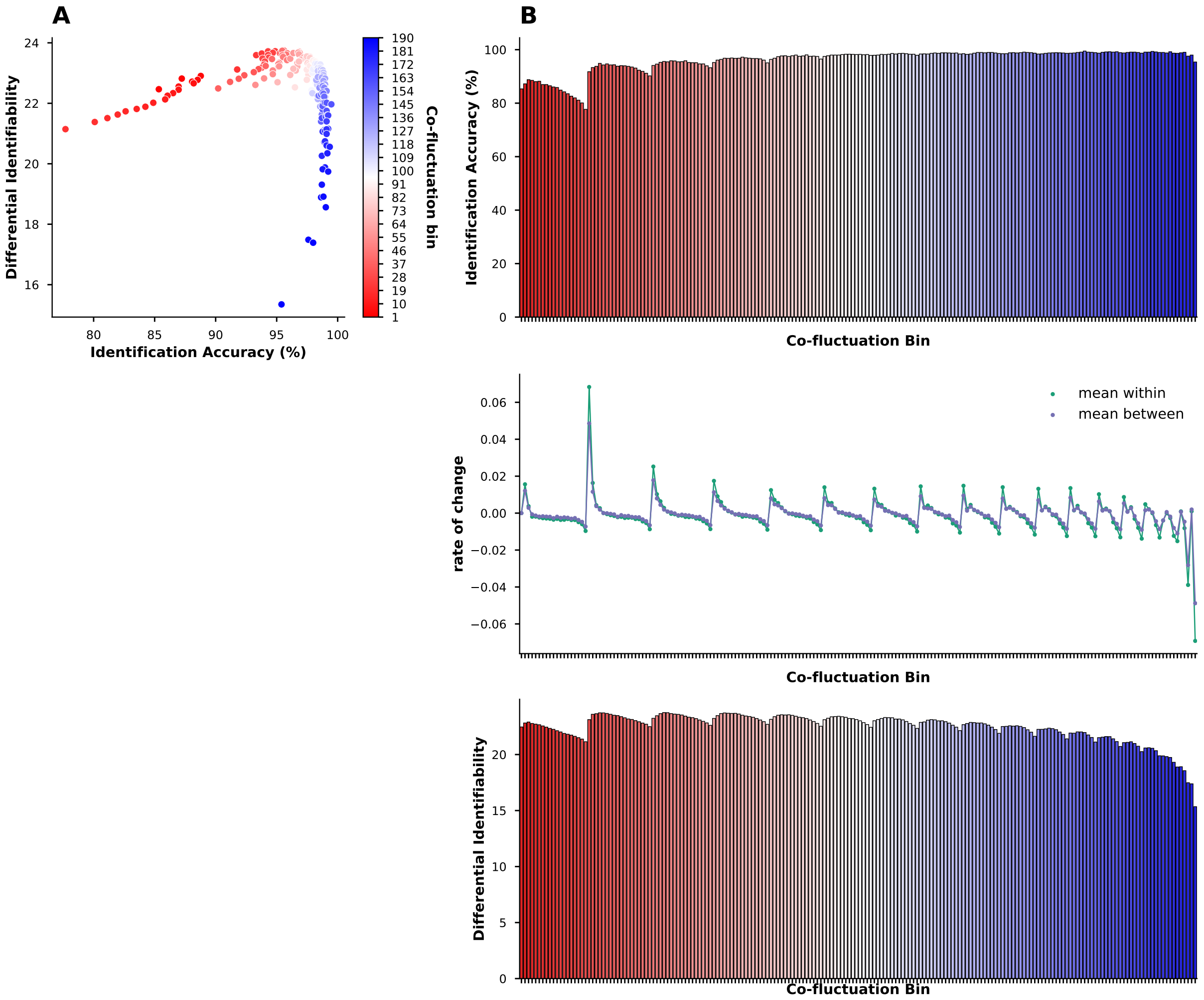


**Figure S2.** Identification accuracy and differential identifiability in the HCP-YA sample using the combined bins strategy: **A)** shows optimal subject specificity of intermediate bins in a scatterplot of identification accuracy and differential identifiability. **B** shows identification accuracy (top row) and differential identifiability (bottom row) as well as the derivative (i.e. difference between one bin and the next) of mean within- and mean-between subject correlations (middle row).

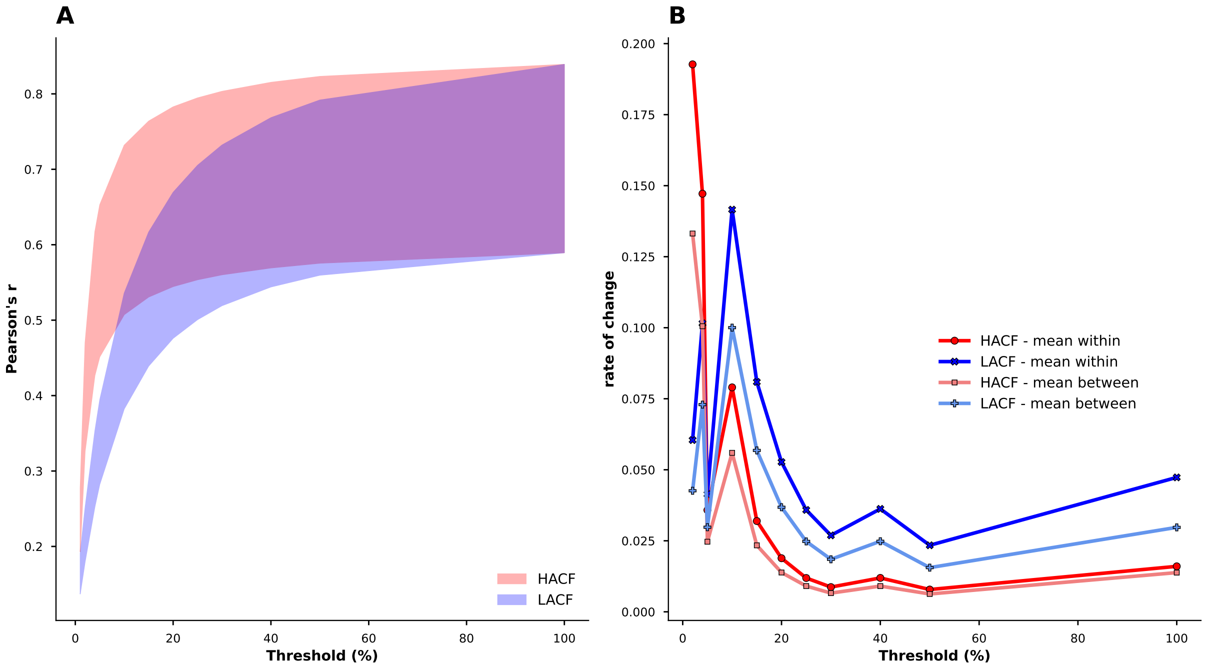


**Figure S3. A)**: Within- and between-subject correlations in the sequential sampling strategy: fill colours indicate co-fluctuation level (i.e. “HACF” or “LACF”). The lower boundary of fill colour shows the mean between-subject correlation, whereas the upper boundary shows the mean within-subject correlation. **B)**: Derivative for each co-fluctuation level’s mean within- and between-subject correlations (i.e. rate of change from one threshold to the next).

**Table S1.** Behavioural prediction targets from the **HCP-YA** sample

| Name | HCP-YA field | Category |
|---|---|---|
| Visual Episodic Memory | PicSeq_Unadj | Cognition |
| Cognitive flexbility (DCCS) | CardSort_Unadj | Cognition |
| Inhibition (Flanker Task) | Flanker_Unadj | Cognition |
| Fluid Intelligence (PMAT) | PMAT24_A_CR | Cognition |
| Reading (pronounciation) | ReadEng_Unadj | Cognition |
| Vocabulary (picture matching) | PicVocab_Unadj | Cognition |
| Processing Speed | ProcSpeed_Unadj | Cognition |
| Delay Discounting | DDisc_AUC_40K | Cognition |
| Spatial Orientation | VSPLOT_TC | Cognition |
| Sustained Attention - Sens. | SCPT_SEN | Cognition |
| Sustained Attention - Spec. | SCPT_SPEC | Cognition |
| Verbal Episodic Memory | IWRD_TOT | Cognition |
| Working Memory (list sorting) | ListSort_Unadj | Cognition |
| Emotional Face Matching | Emotion_Task_Face_Acc | In-Scanner Task Performance |
| Arithmetic | Language_Task_Math_ Avg_Difficulty_Level | In-Scanner Task Performance |
| Story comprehension | Language_Task_Story_ Avg_Difficulty_Level | In-Scanner Task Performance |
| Relational processing | Relational_Task_Acc | In-Scanner Task Performance |
| Social Cognition - random | Social_Task_Perc_Random | In-Scanner Task Performance |
| Social Cognition - interaction | Social_Task_Perc_TOM | In-Scanner Task Performance |
| Working Memory (n-back) | WM_Task_Acc | In-Scanner Task Performance |
| Argreeableness (NEO) | NEOFAC_A | Personality |
| Openness (NEO) | NEOFAC_O | Personality |
| Conscientiousness (NEO) | NEOFAC_C | Personality |
| Neuroticism (NEO) | NEOFAC_N | Personality |
| Extraversion (NEO) | NEOFAC_E | Personality |

**Table S2.** Behavioural prediction targets from the **HCP-A** sample

| Name | NDA data structure | HCP-A field | category |
|---|---|---|---|
| Crystal. Cog. Comp. Score | cogcomp01 | nih_crycogcomp_unadjusted | Cognition |
| Lang./Vocab. Comprehension | tpvt01 | tpvt_uss | Cognition |
| Cog. Flexibility | dccs01 | nih_dccs_unadjusted | Cognition |
| Fluid Cog. Comp. Score | cogcomp01 | nih_fluidcogcomp_unadjusted | Cognition |

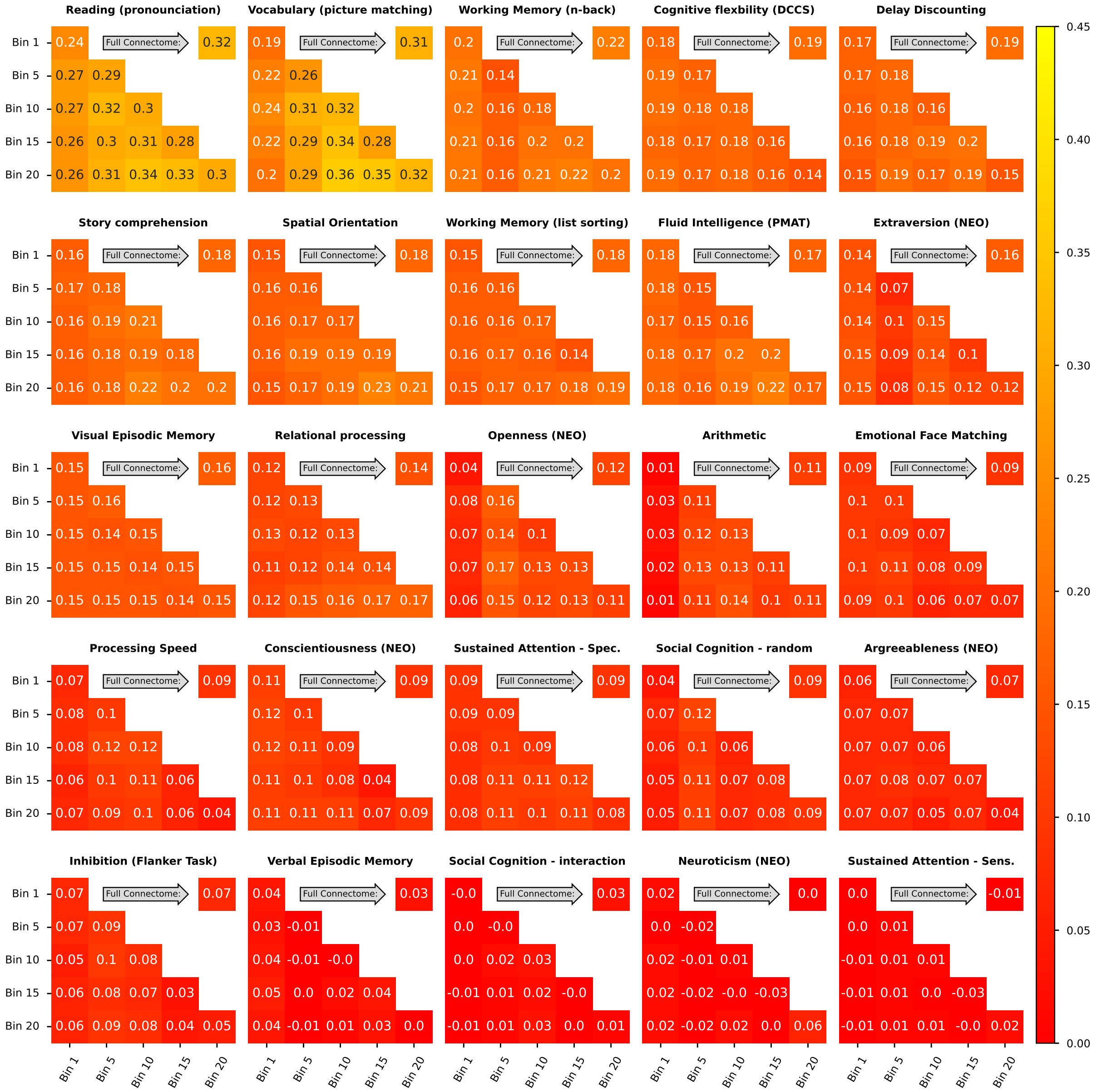

**Figure S4.** Prediction scores (**Pearson's r** between observed and predicted values) in the **HCP-YA** sample for all targets using **kernel ridge regression** averaged across the ten folds in the grouped cross-validation scheme when using combined and individual bins sampling strategies and the **200 area Schaefer parcellation with global signal regression**. Scores for individual bins are displayed on the diagonal, for combined bins off the diagonal. Scores for the full FC using the whole time series are always displayed in the upper right corner.

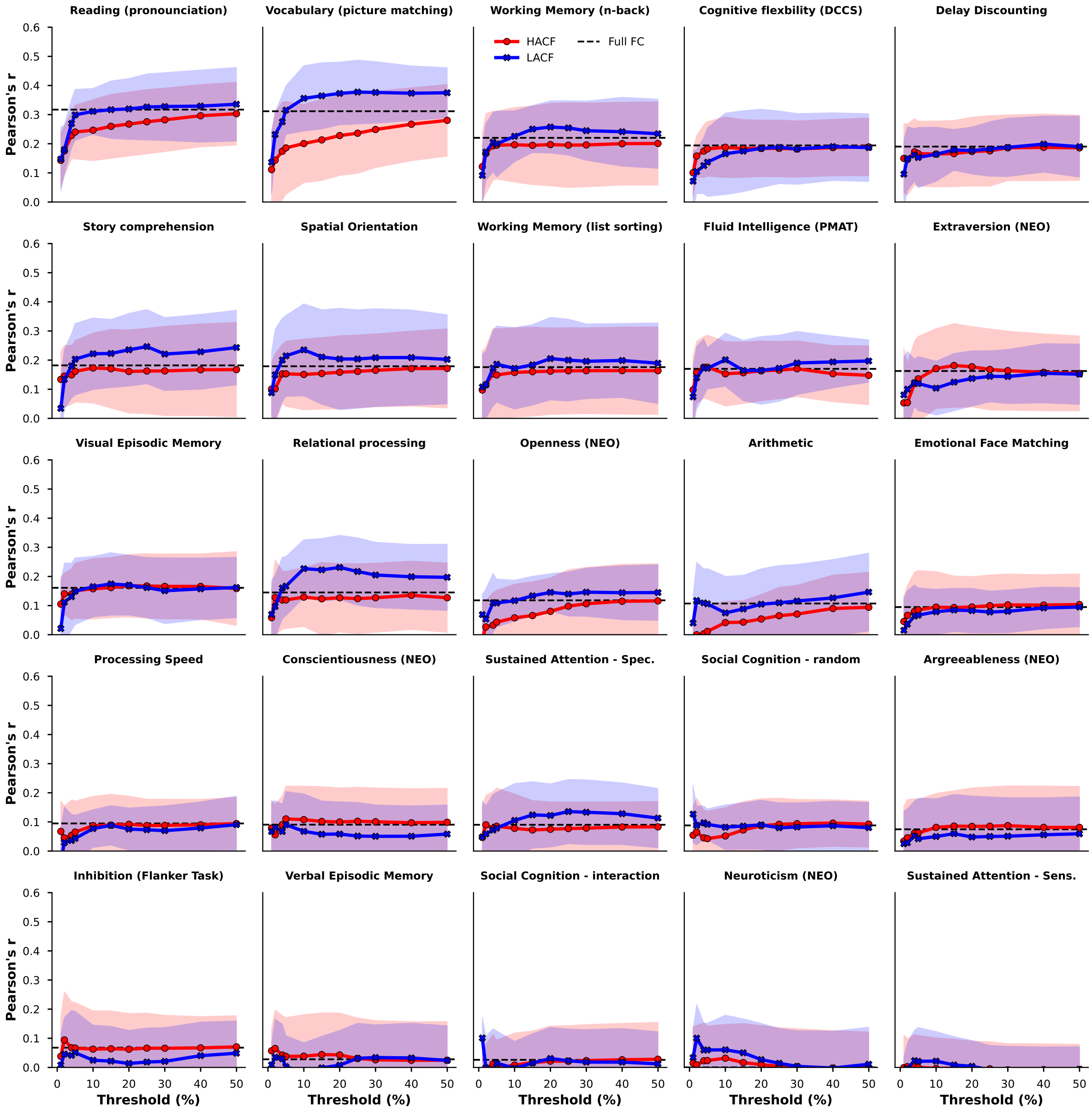


**Figure S5.** Prediction scores (**Pearson's r** between observed and predicted values) in the **HCP-YA** sample for **kernel ridge regression** averaged across the ten folds in the grouped cross-validation scheme when using the **200 area Schaefer parcellation with global signal regression** and FC estimates derived from timepoints at different levels of co-fluctuation magnitude in the sequential sampling strategy.

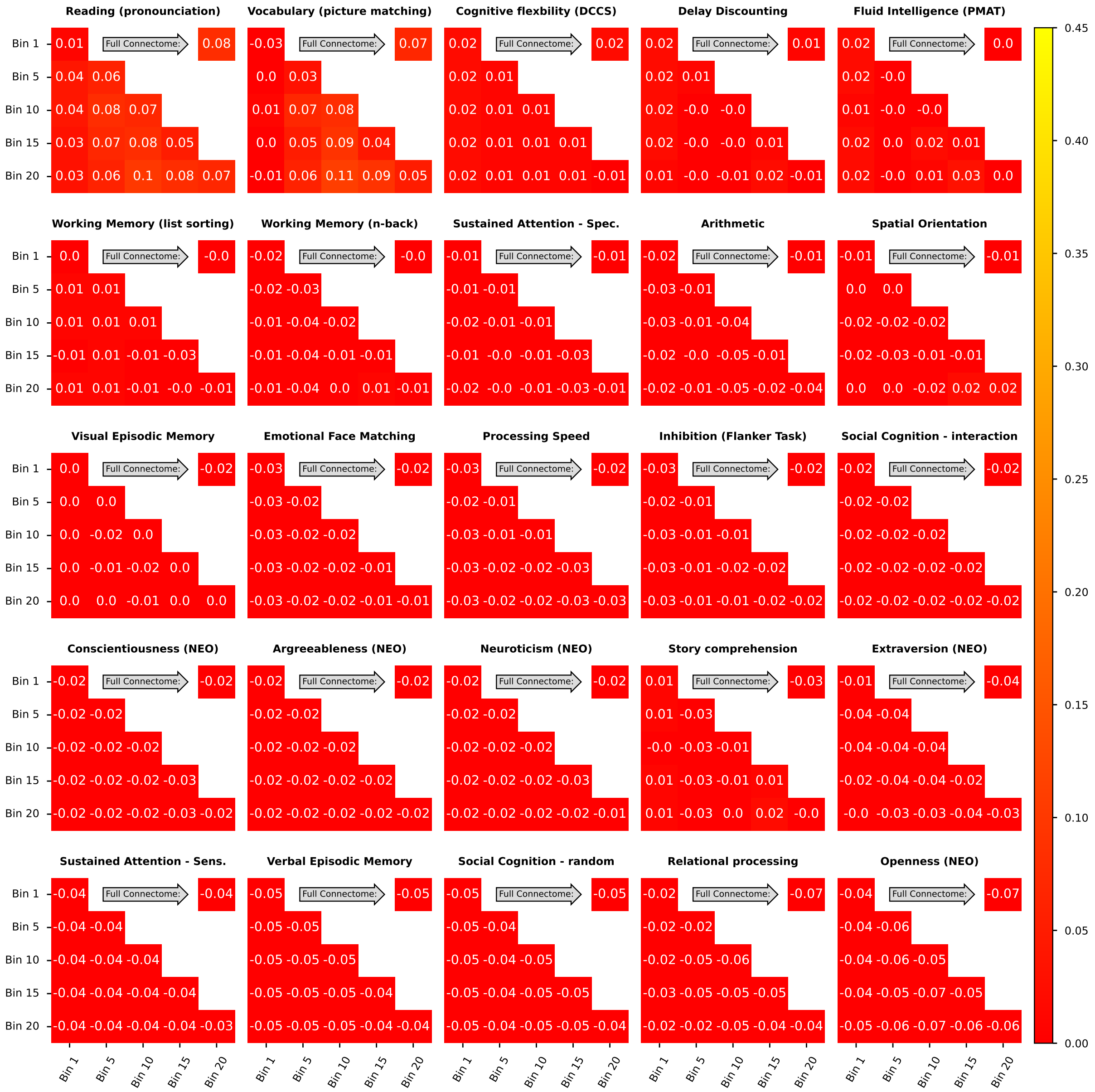


**Figure S6.** Prediction scores (**r-squared**) in the **HCP-YA** sample for all targets using **kernel ridge regression** averaged across the ten folds in the grouped cross-validation scheme when using combined and individual bins sampling strategies and the **200 area Schaefer parcellation with global signal regression**. Scores for individual bins are displayed on the diagonal, for combined bins off the diagonal. Scores for the full FC using the whole time series are always displayed in the upper right corner.

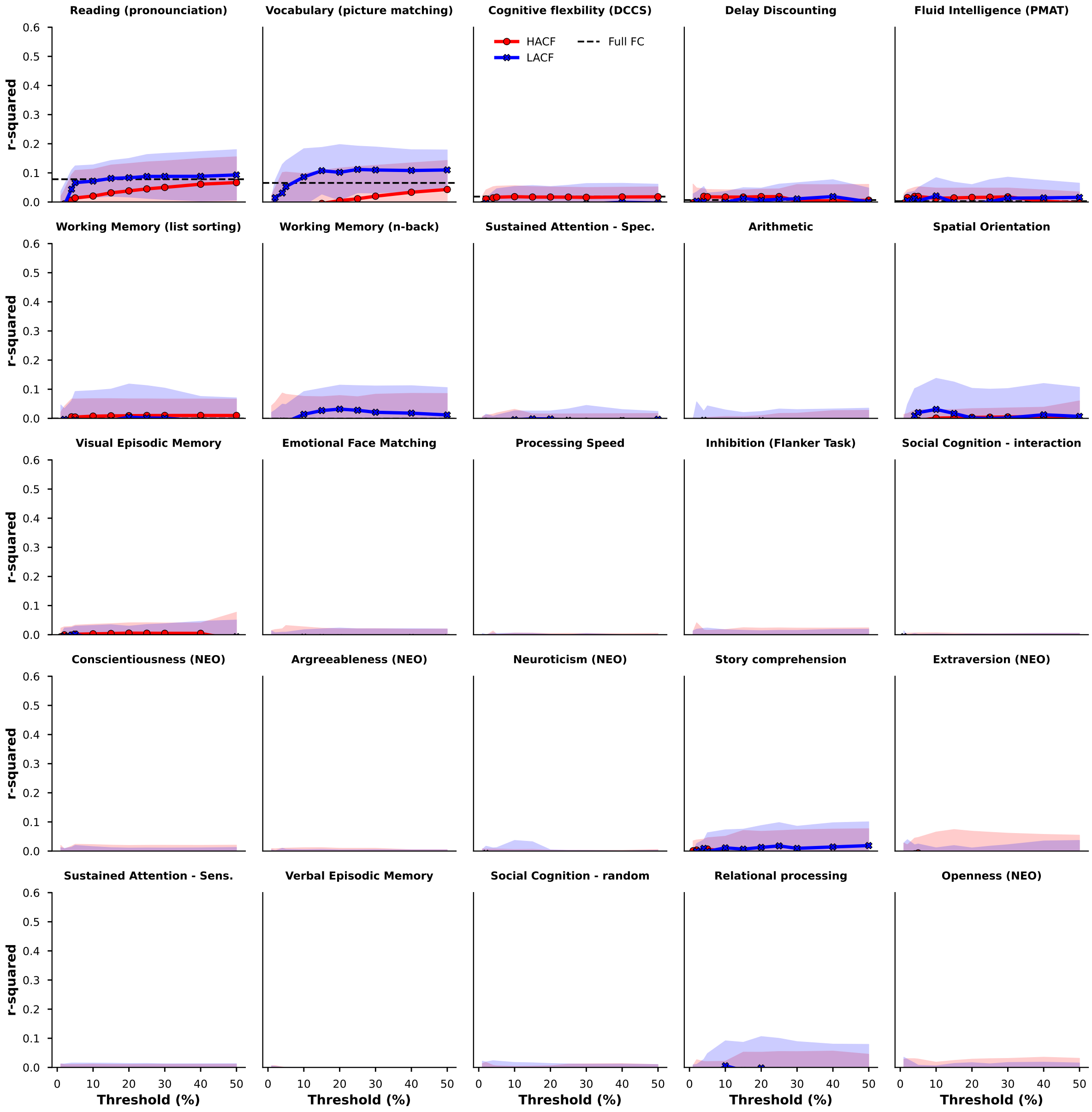


**Figure S7.** Prediction scores (**r-squared** between observed and predicted values) in the **HCP-YA** sample for **kernel ridge regression** averaged across the ten folds in the grouped cross-validation scheme when using the **200 area Schaefer parcellation with global signal regression** and FC estimates derived from timepoints at different levels of co-fluctuation magnitude in the sequential sampling strategy.

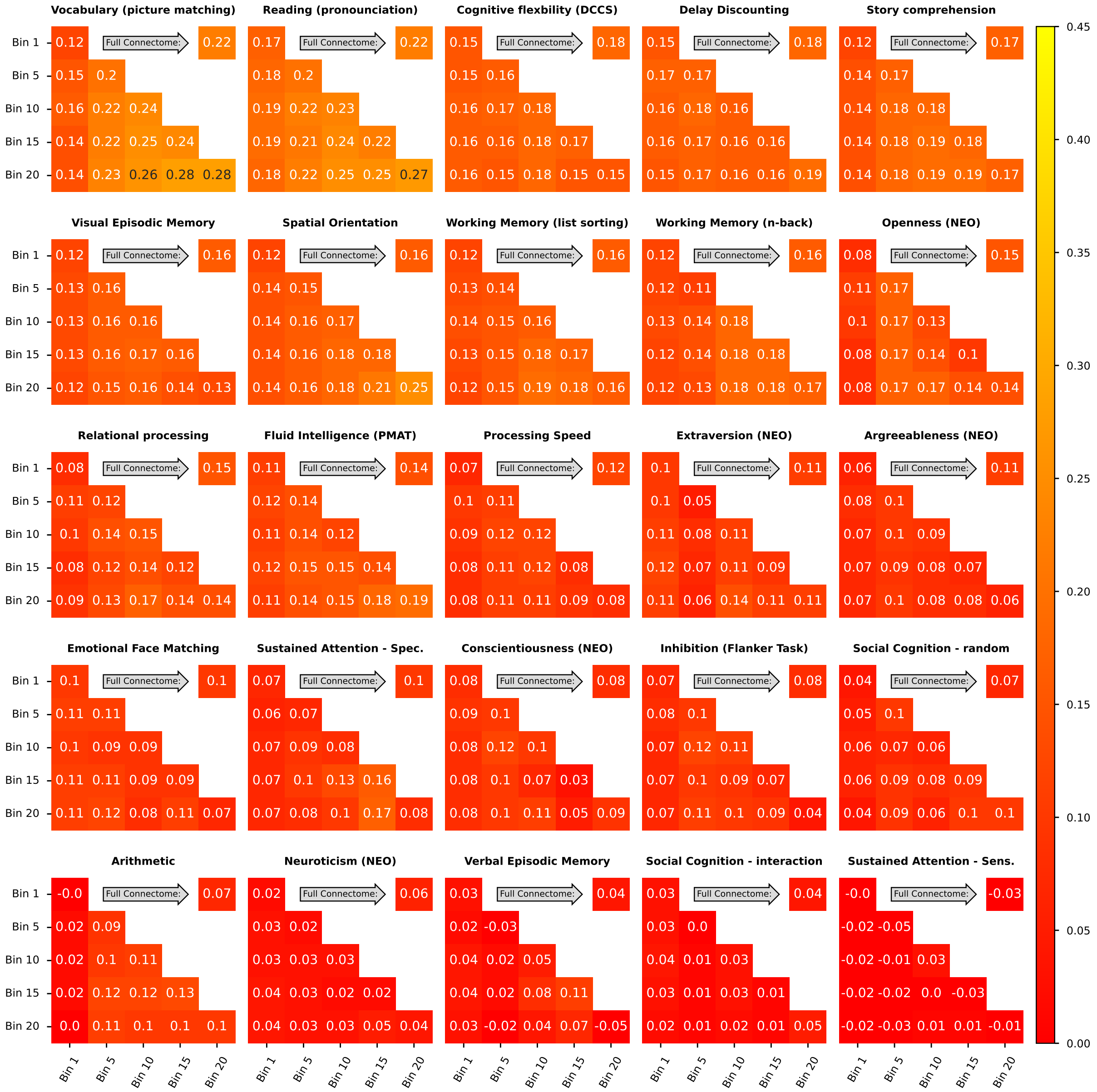

**Figure S8.** Prediction scores (**Pearson's r** between observed and predicted values) in the **HCP-YA** sample for **Connectome-based Predictive Modeling (CBPM)** averaged across the ten folds in the grouped cross-validation scheme when using combined and individual bins sampling strategies and the **200 area Schaefer parcellation**. Scores for individual bins are displayed on the diagonal, for combined bins off the diagonal. Scores for the full FC using the whole time series are always displayed in the upper right corner.

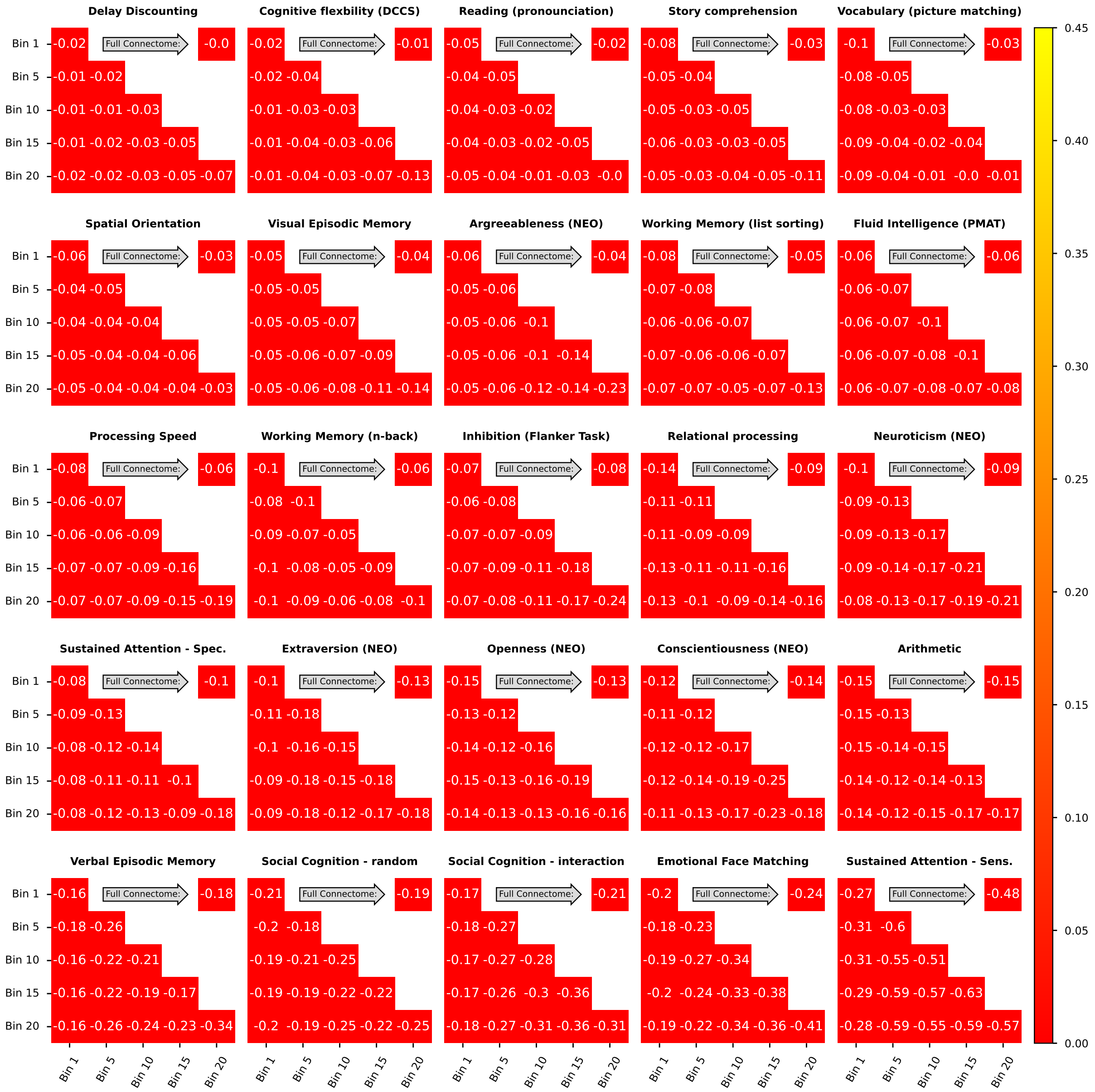


**Figure S9.** Prediction scores (**r-squared**) in the **HCP-YA** sample for **Connectome-based Predictive Modeling (CBPM)** averaged across the ten folds in the grouped cross-validation scheme when using combined and individual bins sampling strategies and the **200 area Schaefer parcellation**. Scores for individual bins are displayed on the diagonal, for combined bins off the diagonal. Scores for the full FC using the whole time series are always displayed in the upper right corner.

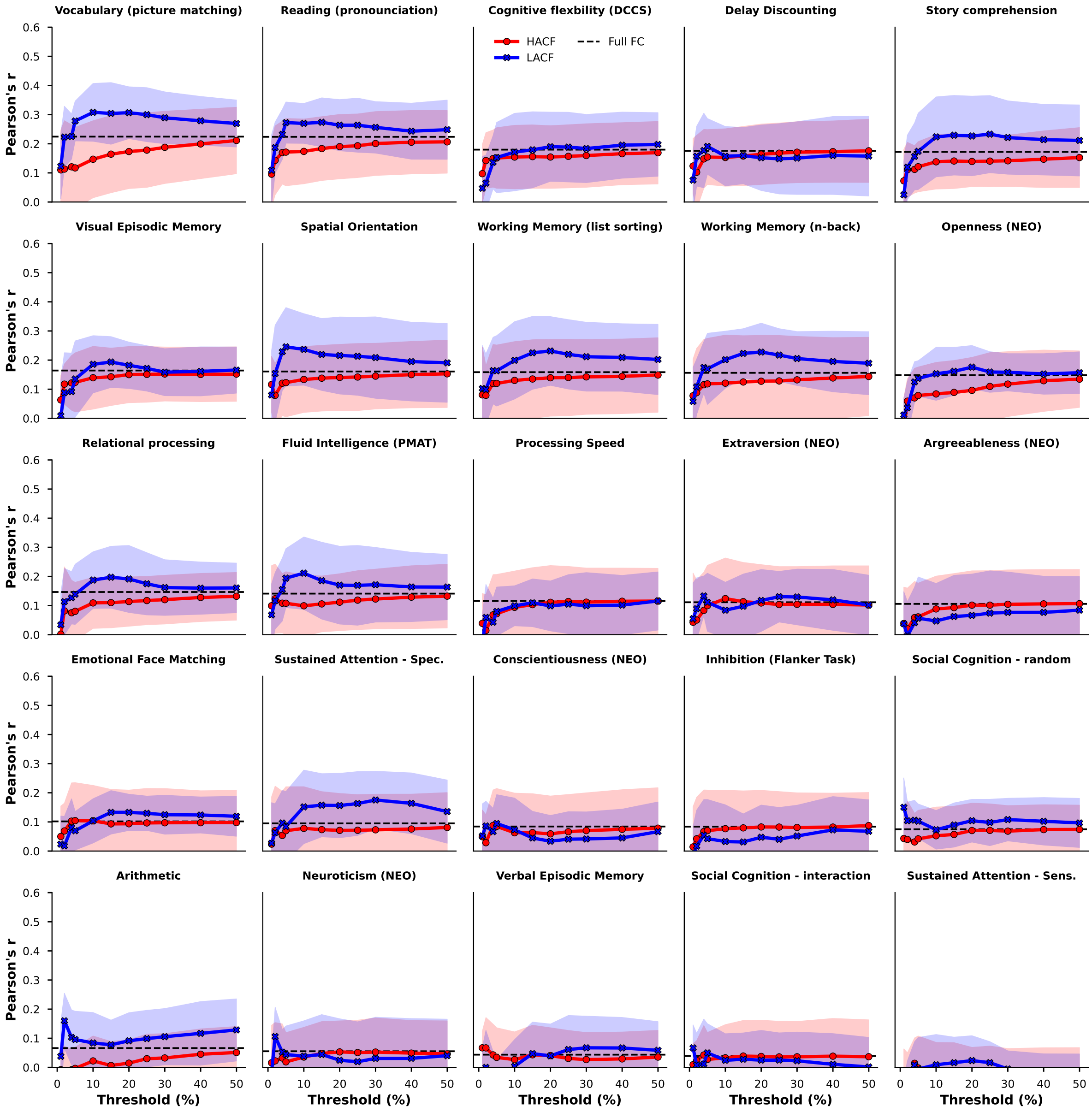


**Figure S10.** Prediction scores (**Pearson's r** between observed and predicted values) in the **HCP-YA** sample for **CBPM** averaged across the ten folds in the grouped cross-validation scheme when using the **200 area Schaefer parcellation** and FC estimates derived from timepoints at different levels of co-fluctuation magnitude in the sequential sampling strategy.

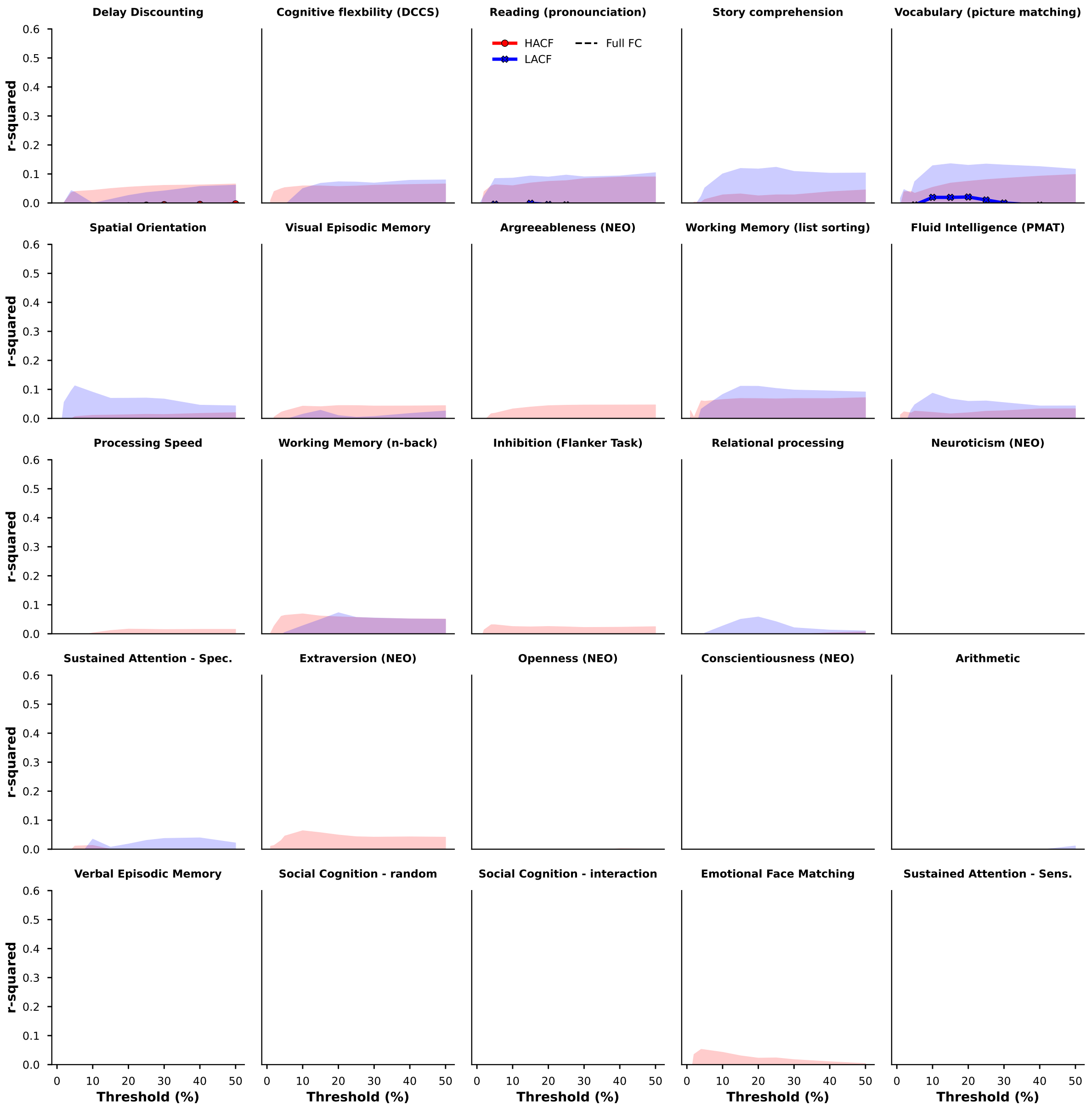


**Figure S11.** Prediction scores (**r-squared**) in the **HCP-YA** sample for **CBPM** averaged across the ten folds in the grouped cross-validation scheme when using the **200 area Schaefer parcellation** and FC estimates derived from timepoints at different levels of co-fluctuation magnitude in the sequential sampling strategy.

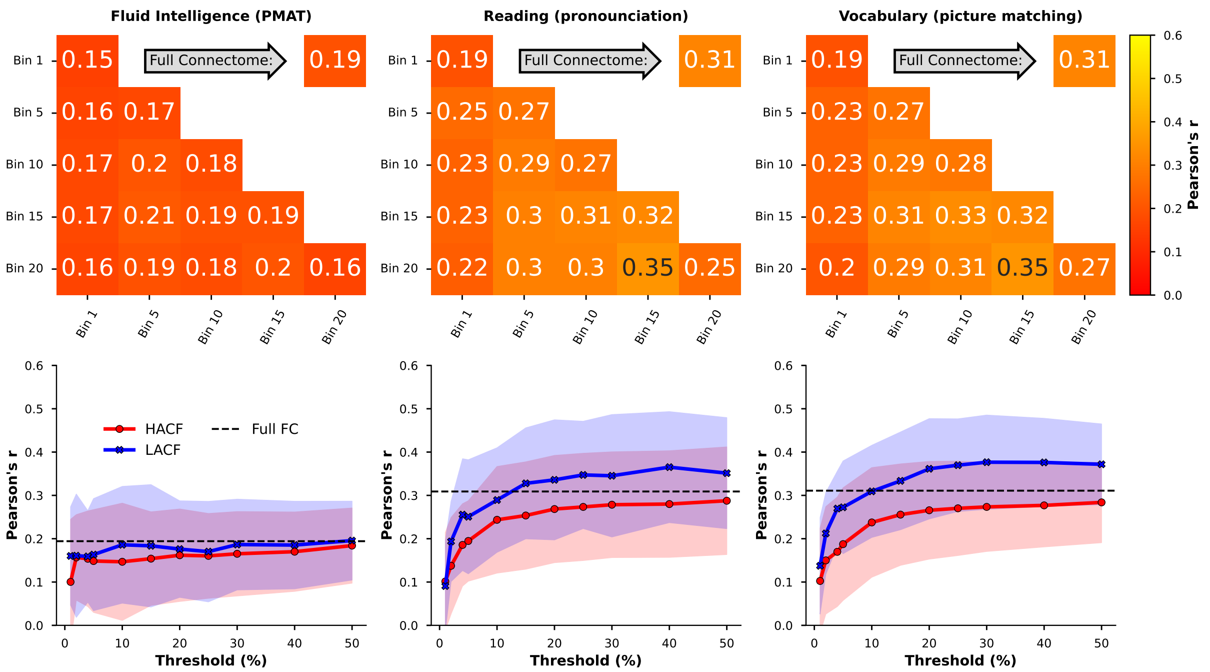


**Figure S12.** Prediction scores (**Pearson's r** between observed and predicted values) in the **HCP-YA** sample for **kernel ridge regression** averaged across the ten folds in the grouped cross-validation scheme when using combined and individual bins sampling strategies and the **200 area Schaefer parcellation without global signal regression**

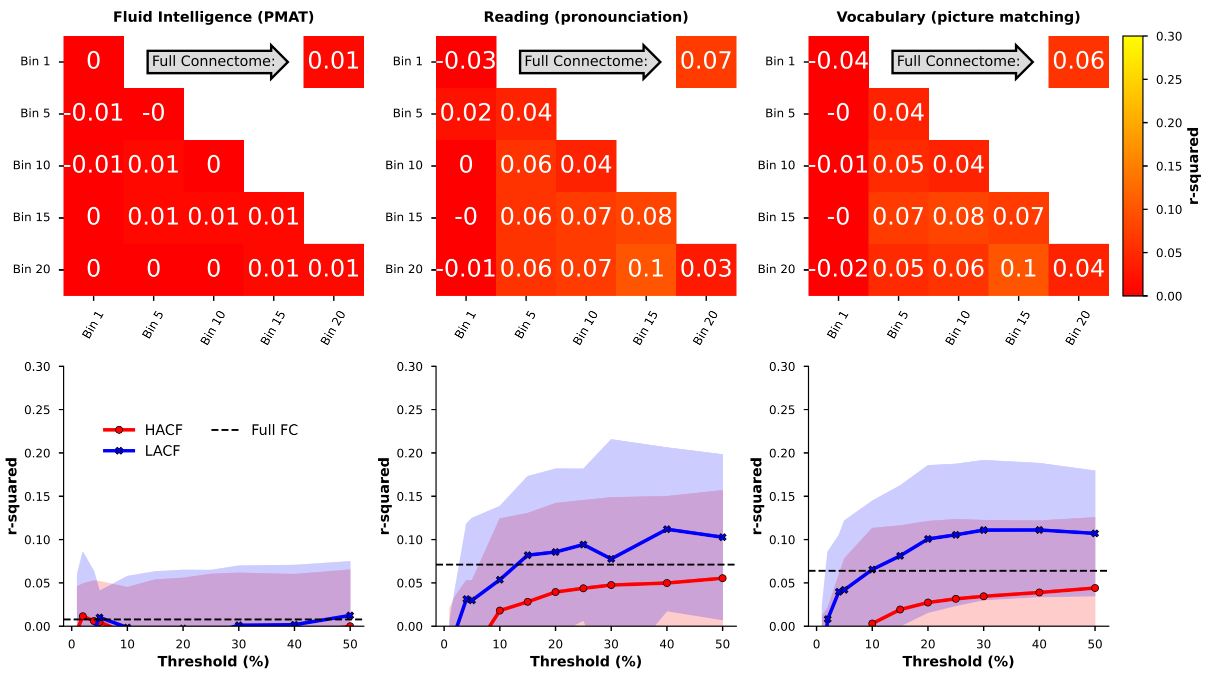


**Figure S13.** Prediction scores (**r-squared** between observed and predicted values) in the **HCP-YA** sample for **kernel ridge regression** averaged across the ten folds in the grouped cross-validation scheme when using combined and individual bins sampling strategies and the **200 area Schaefer parcellation without global signal regression**

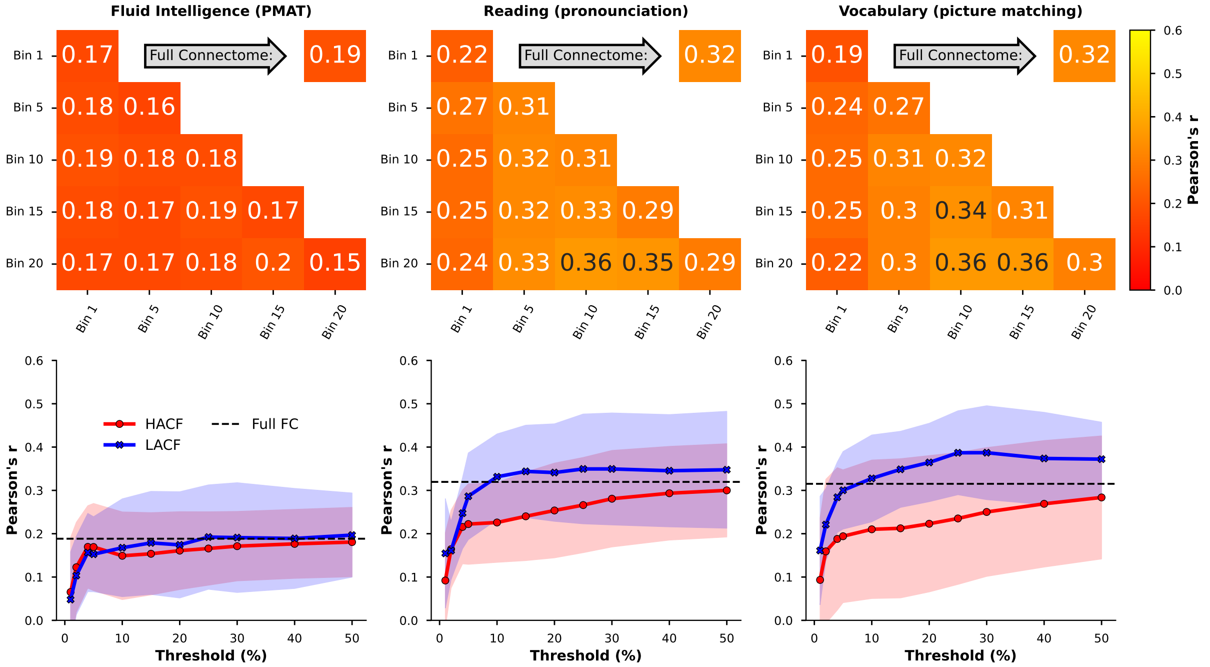


**Figure S14.** Prediction scores (**Pearson's r** between observed and predicted values) in the **HCP-YA** sample for **kernel ridge regression** averaged across the ten folds in the grouped cross-validation scheme when using combined and individual bins sampling strategies and the **300 area Schaefer parcellation with global signal regression**

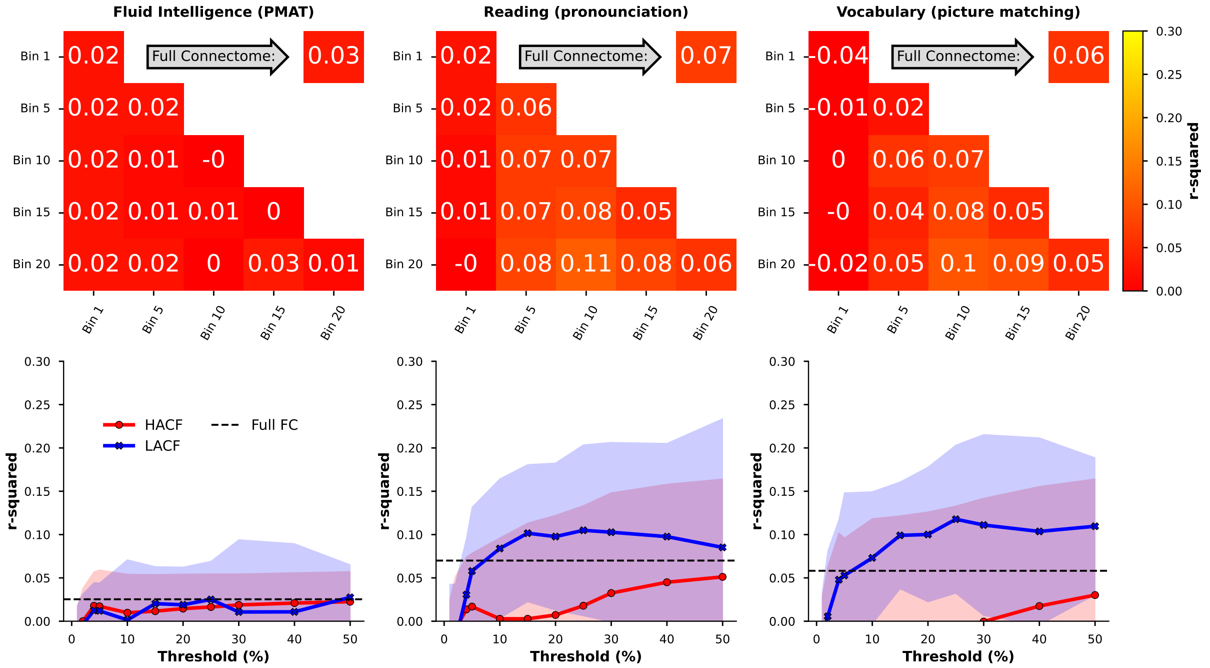


**Figure S15.** Prediction scores (**r-squared** between observed and predicted values) in the **HCP-YA** sample for **kernel ridge regression** averaged across the ten folds in the grouped cross-validation scheme when using combined and individual bins sampling strategies and the **300 area Schaefer parcellation with global signal regression**

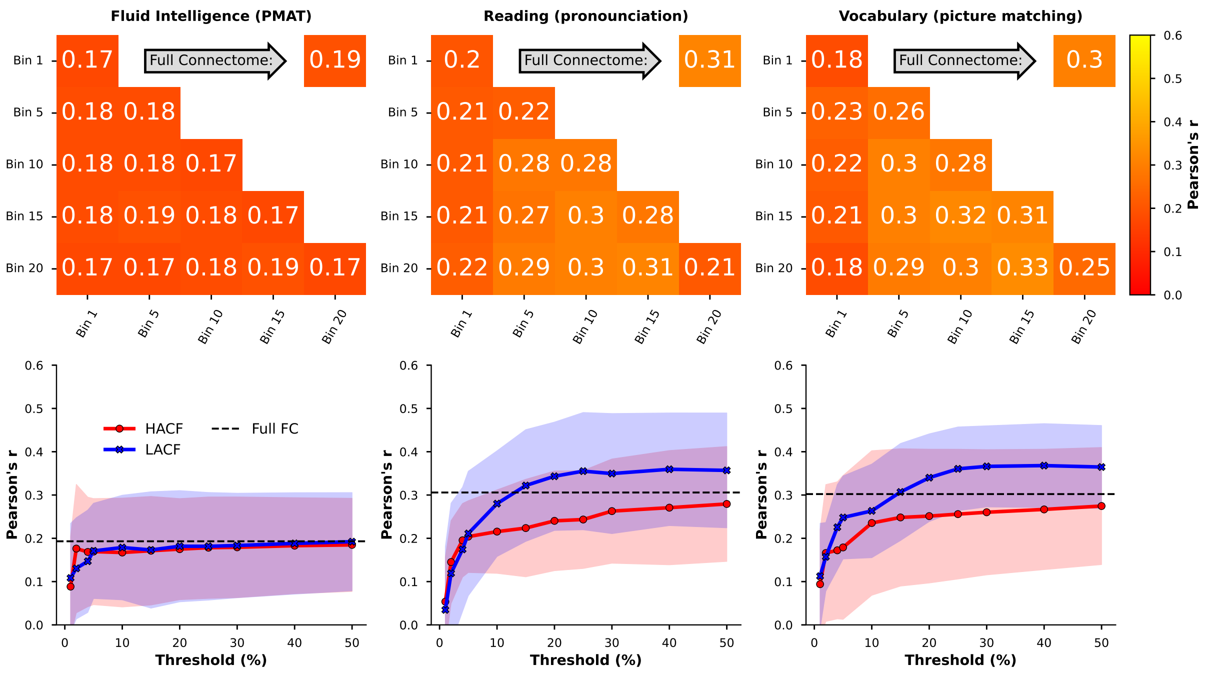


**Figure S16.** Prediction scores (**Pearson's r** between observed and predicted values) in the **HCP-YA** sample for **kernel ridge regression** averaged across the ten folds in the grouped cross-validation scheme when using combined and individual bins sampling strategies and the **300 area Schaefer parcellation without global signal regression**

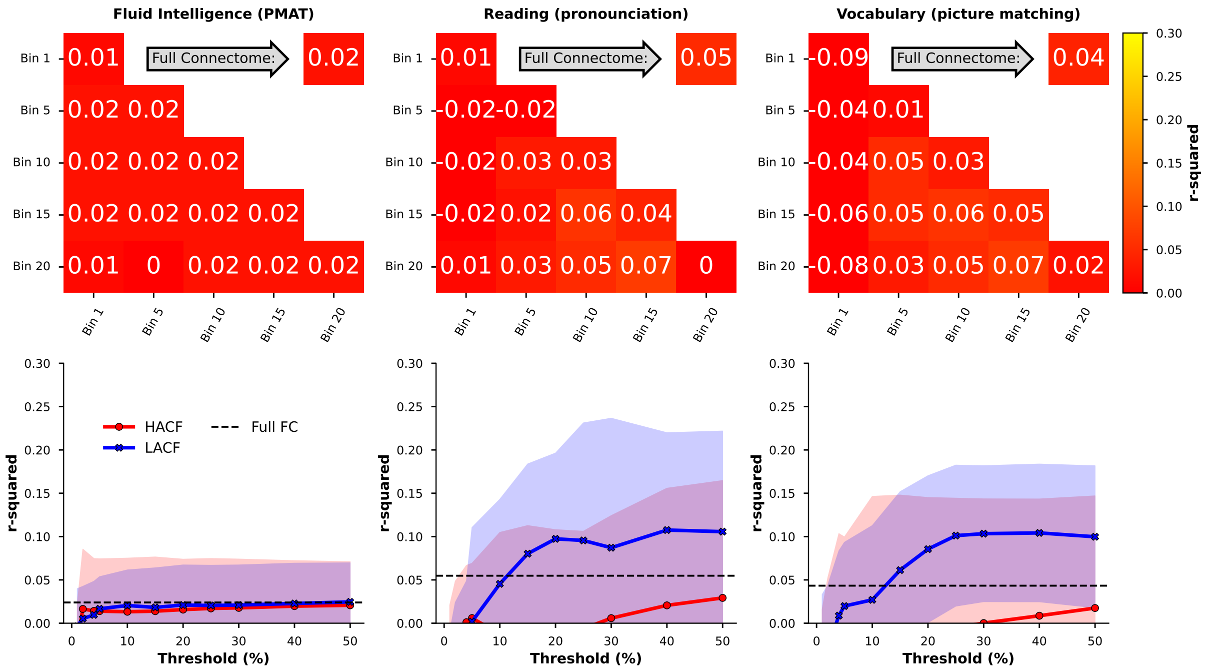


**Figure S17.** Prediction scores (**r-squared** between observed and predicted values) in the **HCP-YA** sample for **kernel ridge regression** averaged across the ten folds in the grouped cross-validation scheme when using combined and individual bins sampling strategies and the **300 area Schaefer parcellation without global signal regression**

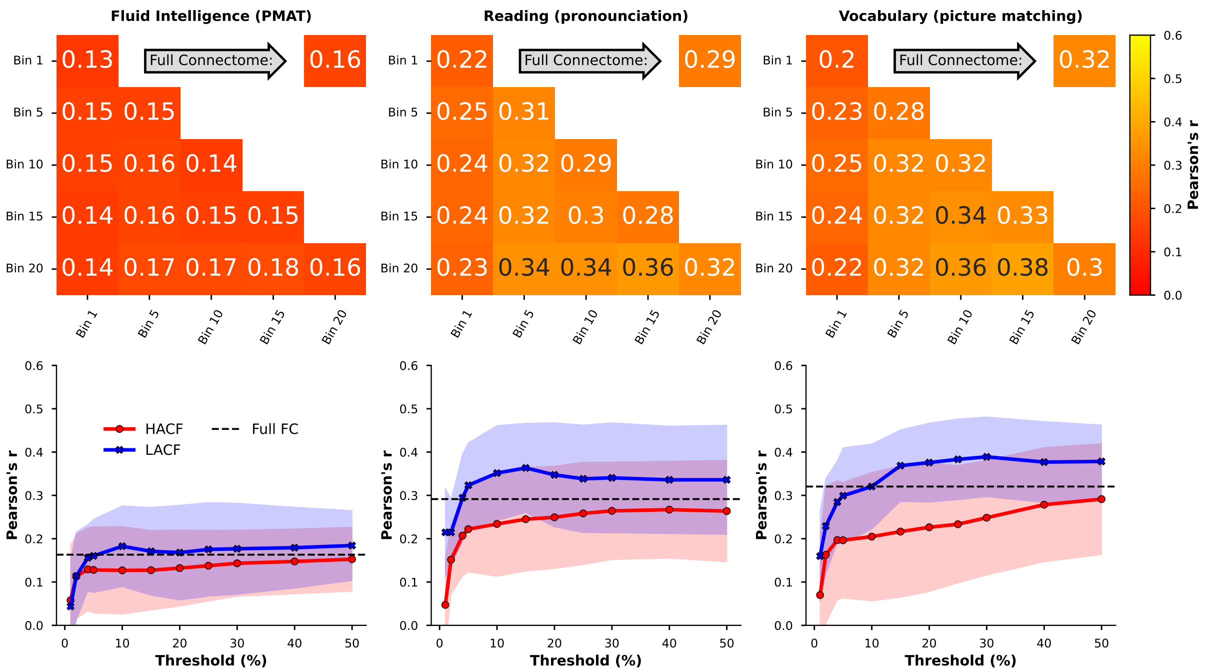


**Figure S18.** Prediction scores (**Pearson's r** between observed and predicted values) in the **HCP-YA** sample for **kernel ridge regression** averaged across the ten folds in the grouped cross-validation scheme when using combined and individual bins sampling strategies and the **400 area Schaefer parcellation with global signal regression**

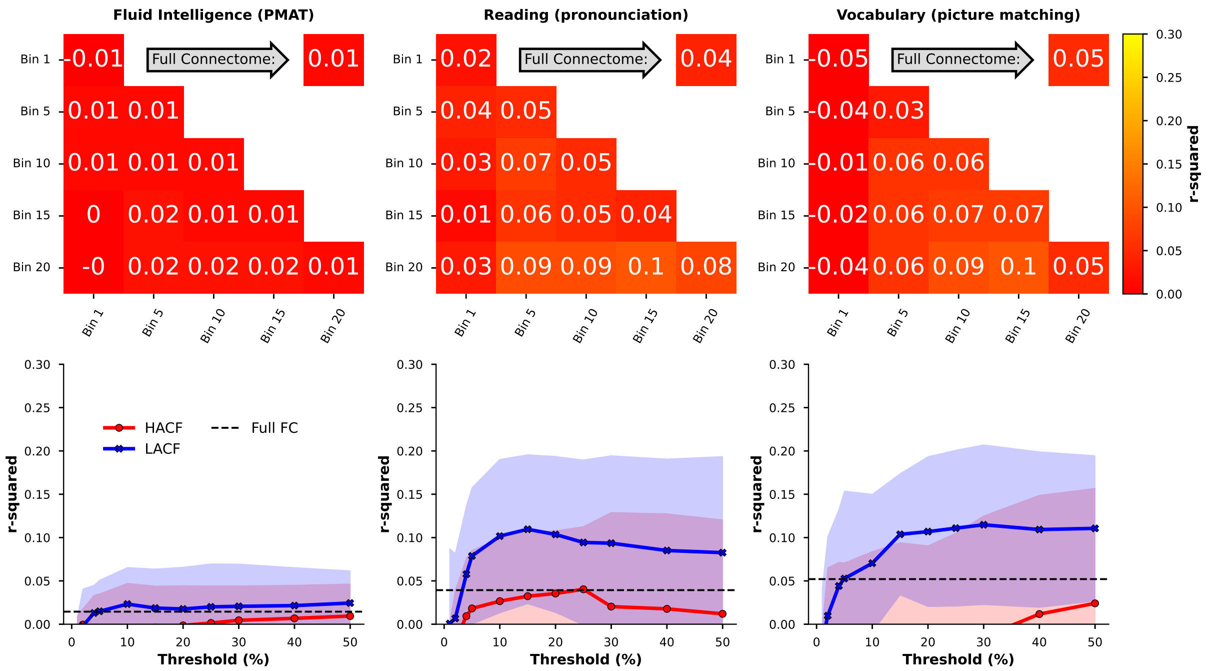


**Figure S19.** Prediction scores (**r-squared** between observed and predicted values) in the **HCP-YA** sample for **kernel ridge regression** averaged across the ten folds in the grouped cross-validation scheme when using combined and individual bins sampling strategies and the **400 area Schaefer parcellation with global signal regression**

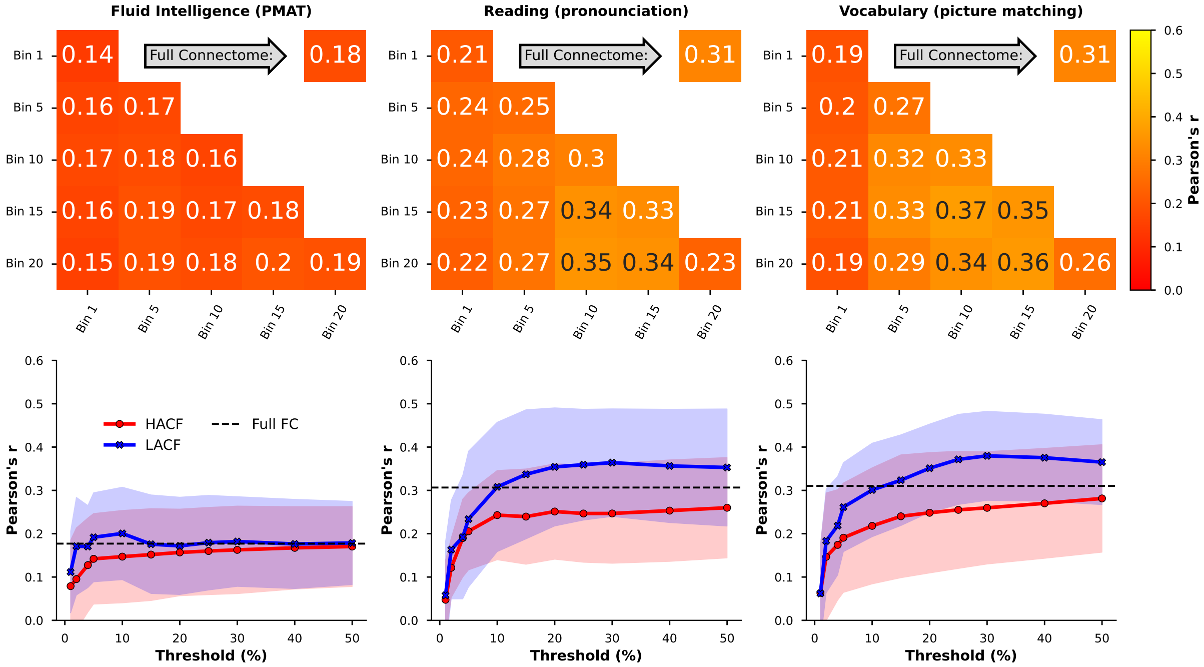


**Figure S20.** Prediction scores (**Pearson's r** between observed and predicted values) in the **HCP-YA** sample for **kernel ridge regression** averaged across the ten folds in the grouped cross-validation scheme when using combined and individual bins sampling strategies and the **400 area Schaefer parcellation without global signal regression**

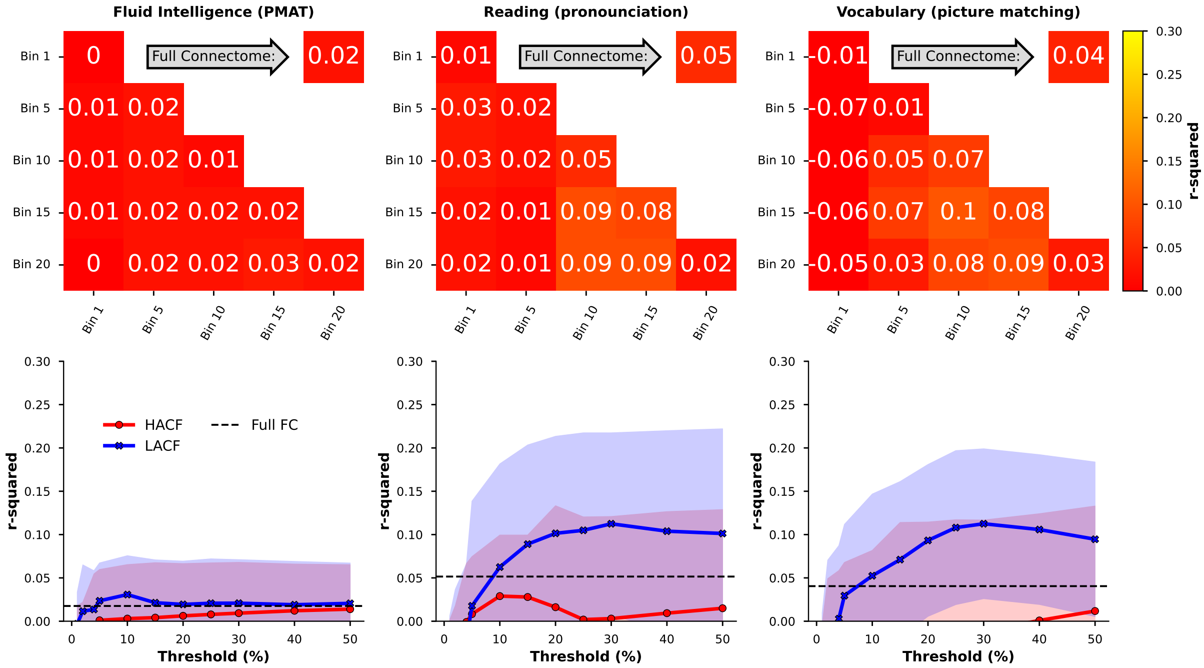


**Figure S21.** Prediction scores (**r-squared** between observed and predicted values) in the **HCP-YA** sample for **kernel ridge regression** averaged across the ten folds in the grouped cross-validation scheme when using combined and individual bins sampling strategies and the **400 area Schaefer parcellation without global signal regression**

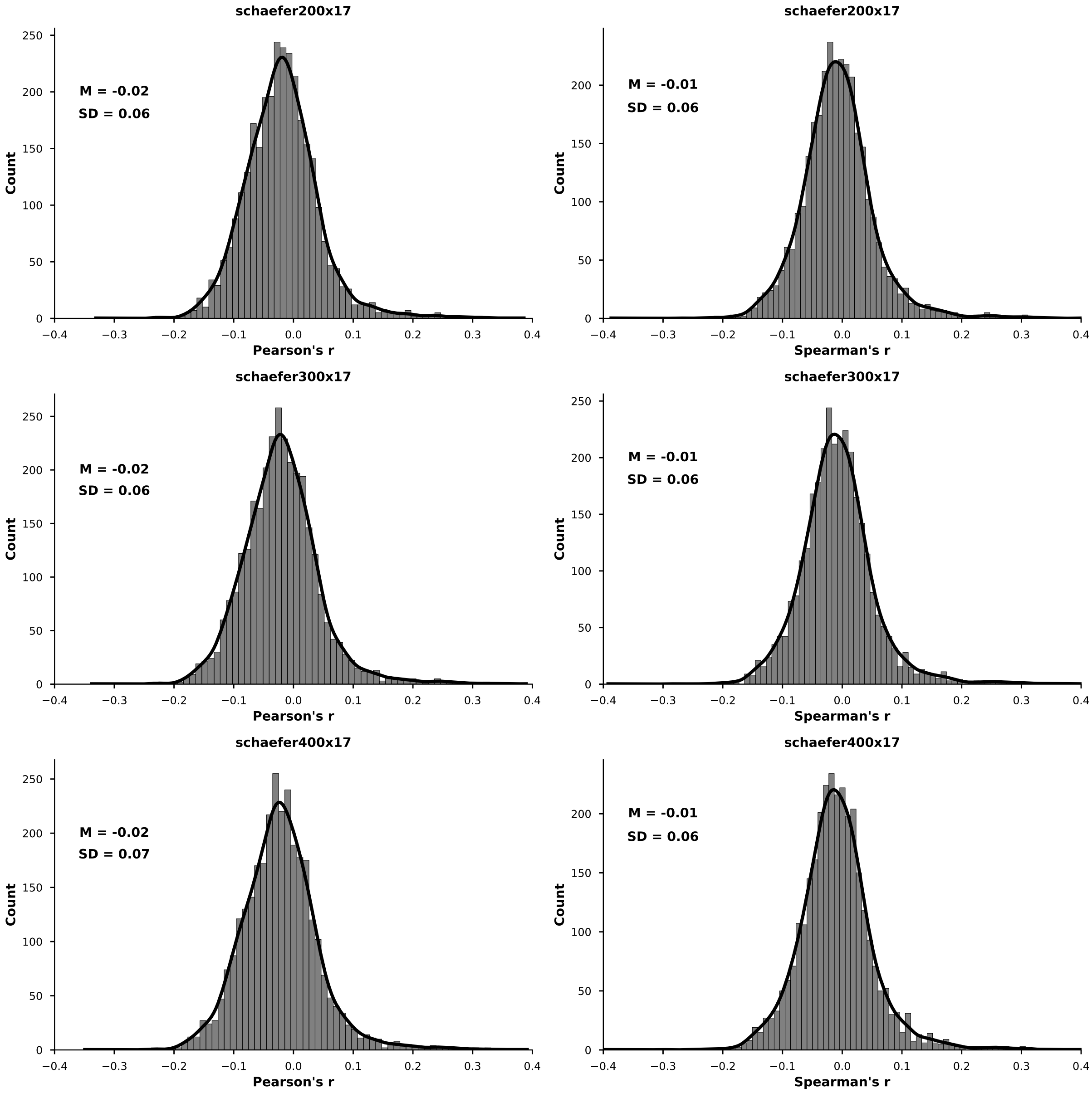


**Figure S22.** Distribution of correlations between RSS and FD for every subject and every rs-fMRI run using three different parcellations and both Spearman's and Pearson's correlation coefficients.

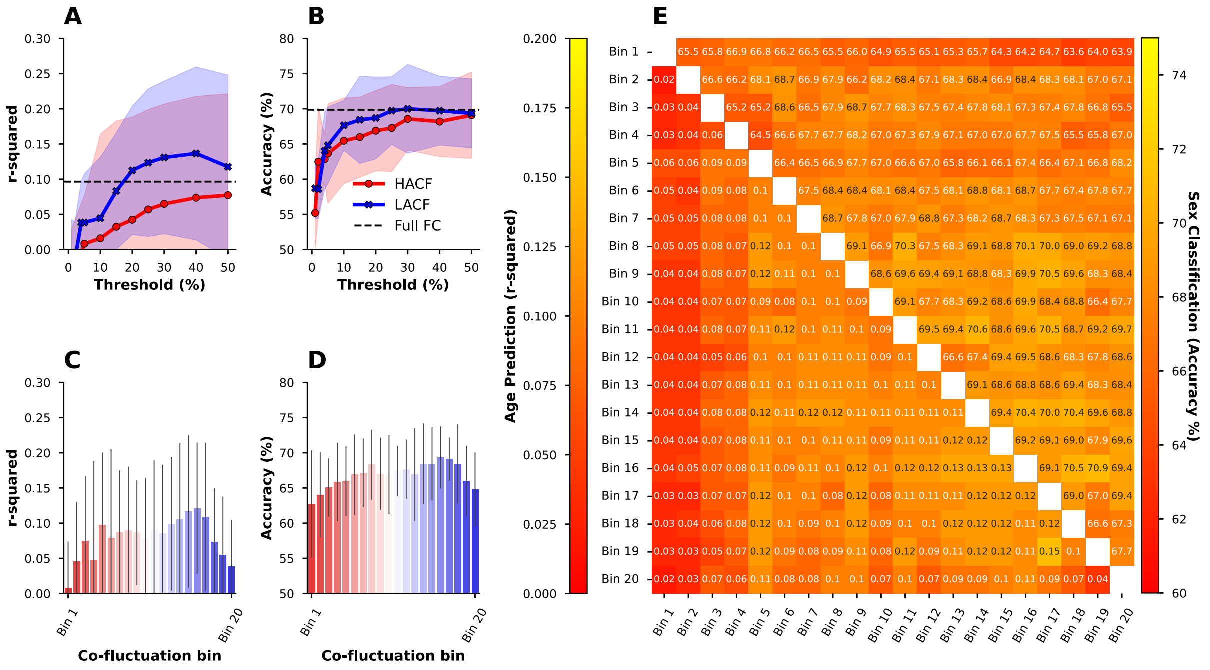


**Figure S23.** Age prediction scores (**r-squared**) and sex classification accuracy in the **HCP-YA** sample for the sequential (**A** and **B**), individual bins (**C** and **D**) and the combined bins (**E**) sampling strategies. Here, in sex prediction we use a support vector classifier (SVC) with a linear kernel.

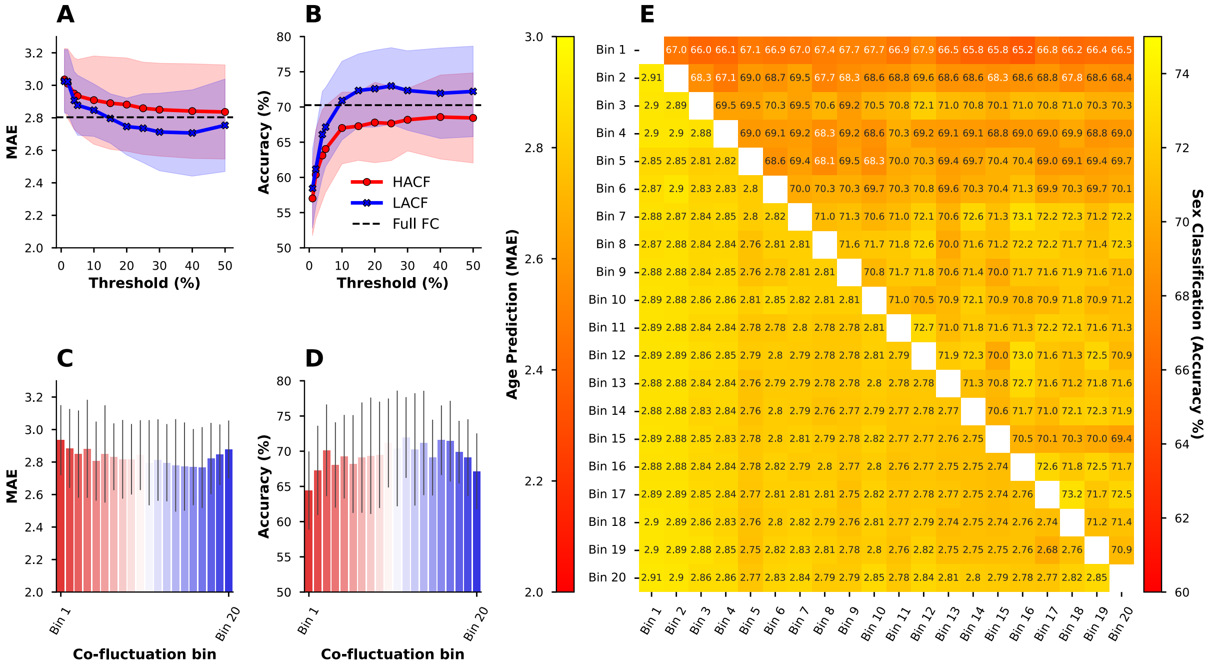


**Figure S24.** Age prediction scores (**MAE**) and sex classification accuracy in the **HCP-YA** sample for the sequential (**A** and **B**), individual bins (**C** and **D**) and the combined bins (**E**) sampling strategies. Here, in sex prediction we use a SVC with a radial basis function (RBF) kernel.

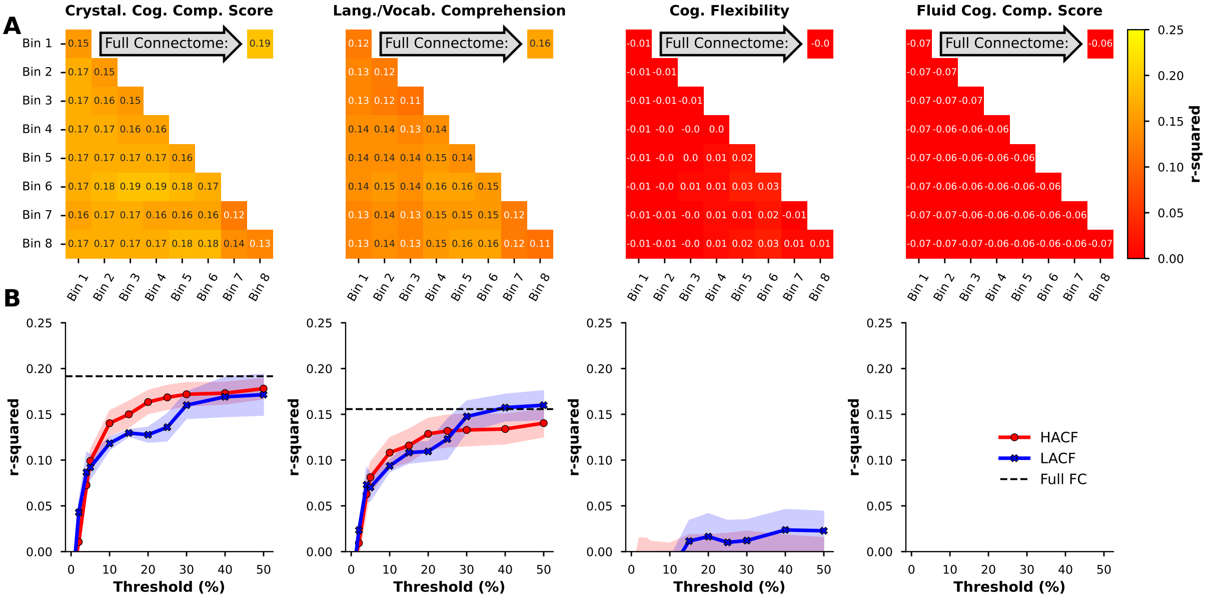


**Figure S25.** Prediction accuracy (**r-squared**) for four cognitive targets in the **HCP-A** sample using the individual and combined bins strategy (**A**; individual bins are on the diagonal) and the sequential sampling strategy (**B**).

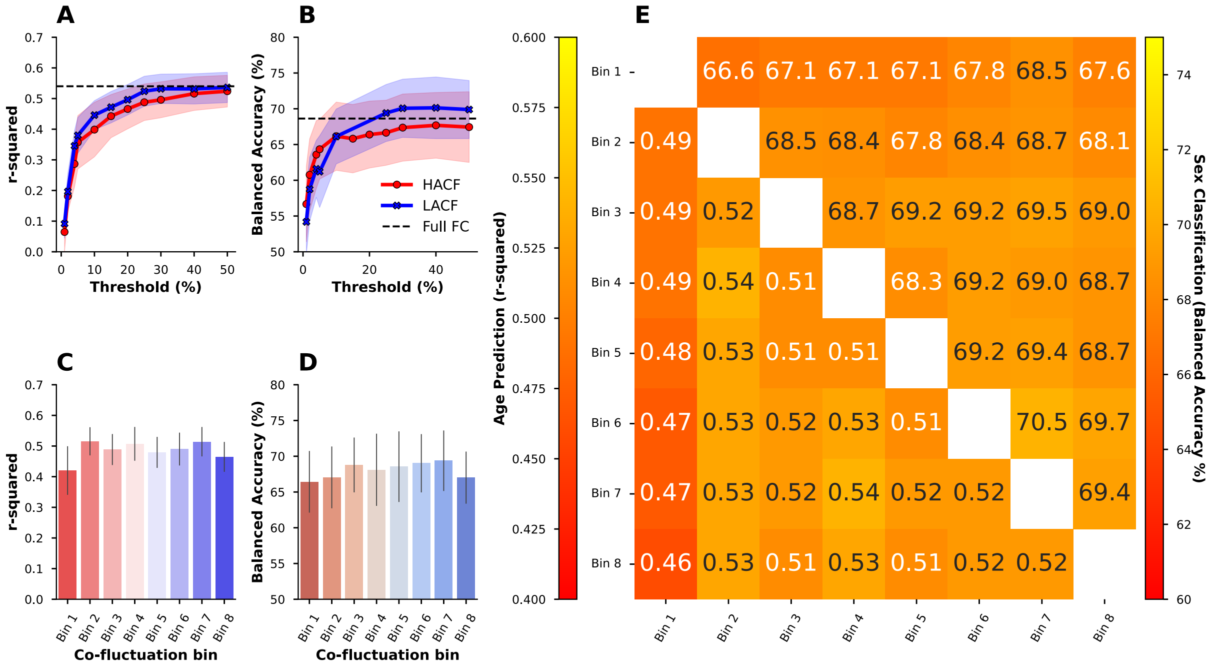


**Figure S26.** Age prediction (**r-squared**) and sex classification (balanced accuracy) in the **HCP-A** sample using the sequential (**A** and **B**), individual bins (**C** and **D**), and combined bins (**E**) sampling strategies using the HCP-A sample. Here, in sex prediction we use a SVC with a linear kernel.

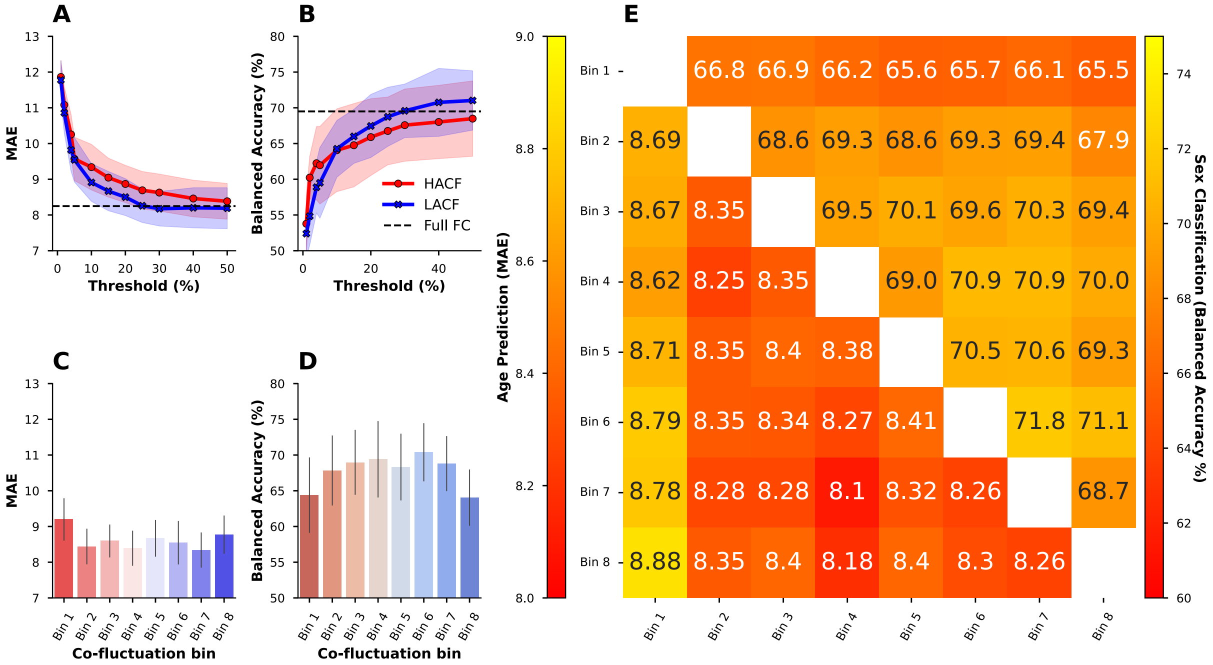


**Figure S27.** Age prediction (**MAE**) and sex classification (balanced accuracy) in the **HCP-A** sample using the sequential (**A** and **B**), individual bins (**C** and **D**), and combined bins (**E**) sampling strategies using the HCP-A sample. Here, in sex prediction we use a SVC with a RBF kernel.
